# Acute DOI exposure drives cortical hyperexcitability and functional network remodeling

**DOI:** 10.64898/2026.05.18.726129

**Authors:** Ido Haber, Ilhan Bok, Benjamin Kutler, Maya Nornberg, Matthew I. Banks, Aviad Hai

## Abstract

Serotonergic psychoplastogens can produce durable cortical remodeling, but how a brief exposure to the 5-HT2A agonist 2,5-dimethoxy-4-iodoamphetamine (DOI) reshapes population activity and functional connectivity remains unclear. We recorded primary rat cortical cultures on spatially defined microelectrode arrays before and after acute DOI exposure using a within-culture repeated-measures design, with a separate ketanserin + DOI arm to probe 5-HT2A receptor involvement. Network activity was summarized from spikes, bursts, and functional connectivity estimated with Pearson cross-correlation and the rate-corrected spike-time tiling coefficient. Following DOI exposure, mean firing rate increased across all six wells, burst timing accelerated, and functional-network metrics showed a convergent but sensitivity-limited shift toward shorter characteristic path length. Ketanserin + DOI exposure reduced population bursting and prolonged inter-burst intervals, while descriptive path-length shortening persisted. Path-length shortening persisted under the rate-corrected STTC estimator, suggesting that the connectivity shift was not simply a firing-rate artifact. These findings show that acute DOI exposure can move dissociated cortical cultures into a hyperexcitable population state with altered functional-network dynamics. Multiplexed cortical network recordings therefore provide a tractable bridge between molecular psychoplastogen biology and systems-level circuit outcomes relevant to durable therapeutic plasticity.

**Highlights:**

- MEA recordings reveal post-acute DOI responses in cortical cultures
- DOI increases spontaneous firing and accelerates burst timing
- DOI shifts functional network structure toward integration
- Ketanserin + DOI suppresses population bursting while preserving path-length shortening

## Introduction

Serotonergic psychedelics commonly referred to as psychoplastogens have re-emerged as a translationally promising class of small molecules that produce rapid, sustained structural and functional remodeling of cortical circuits following a single exposure [1,2,3,4,5]. Compounds in this class — including the classical tryptamines (psilocybin, N,N-dimethyltryptamine), the ergoline lysergic acid diethylamide (LSD), and the substituted phenethylamine 2,5-dimethoxy-4-iodoamphetamine (DOI) — increase dendritic complexity, spinogenesis, and excitatory drive on cortical pyramidal neurons on time-scales that outlast the acute exposure, including effects initiated by transient exposure and expressed after ligand removal [6,7,8,9]. Several recent clinical trials of psilocybin for treatment-resistant depression have lent these observations translational urgency, and the field has converged on the hypothesis that the molecular machinery driving the acute psychedelic experience may also be the substrate for sustained therapeutic plasticity [1,3,4].

Mechanistically, the canonical molecular target for classical psychedelics is the 5-HT2A receptor (5-HT2AR). Agonist binding on cortical pyramidal cells triggers Gq- and β-arrestin-coupled signaling cascades that drive spine enlargement on a minute timescale and dendritic growth over hours to days [2,6,7,10]. DOI is a potent pharmacological agent for probing 5-HT2A function [11]: it has high affinity for 5-HT2A/2C and promotes neuritogenesis, spinogenesis, and synaptogenesis in cortical cultures through TrkB/mTOR-linked mechanisms that are attenuated by ketanserin [6,9]. DOI has also been reported to rapidly engage Trk-related trophic signaling following treatment, including TrkA Tyr490 phosphorylation in SK-N-SH cells and TrkB tyrosine phosphorylation in lymphoblastoid cells [12].

Bridging the molecular and systems levels of description requires an in vitro preparation in which a cultured cortical network is exposed uniformly to controlled pharmacological interventions while activity is sampled from many spatially defined electrodes — bypassing the blood-brain barrier, systemic pharmacokinetics, and hemodynamic confounds that complicate in vivo preparations. Multi-electrode arrays (MEAs) coupled to dissociated cortical cultures have served this role for nearly two decades [13]. Pioneering work established that rat cortical neurons plated on MEAs develop reproducible spontaneous spiking and bursting, that these signatures stabilize across the third week in vitro, and that pharmacological perturbations can be quantified with concentration-response statistics approaching the rigor of conventional electrophysiology [14,15,16]. The preparation has been benchmarked for inter-laboratory reproducibility [17] and has yielded a stable list of network features that survive cross-line and cross-lab comparison [18]. Yet this MEA platform has not been used to characterize serotonergic psychoplastogens at the population-network level. The closest precedent is Varley et al. (2024) [19], who examined information-processing metrics under the psychedelic N,N-dipropyltryptamine in organotypic cortical slices; a characterization of DOI’s network-level effects in dissociated cultures using standard graph-theoretic metrics remains unreported.

The structural plasticity literature has relied on single-cell readouts (Sholl analysis, spine counts, calcium imaging) [6,7,12], and in vivo electrophysiology has probed freely behaving animals but not the isolated cortical network [20,21]. At the single-neuron level, the picture is ambiguous. Hu et al. (2016) [22] showed that DOI *depresses* spontaneous firing in patch-clamped cultured cortical neurons via 5-HT2A/2C activation, while in the same preparation DOI nearly doubled the frequency of miniature excitatory postsynaptic currents (mEPSCs; 3.9 → 7.4 Hz) without changing mEPSC amplitude — a synaptic effect that, considered alone, would favor enhanced spontaneous firing rather than depression. Puig et al. (2003) [23] likewise demonstrated that systemic DOI produced mixed effects across identified mPFC pyramidal neurons, with excitation, inhibition, and no-response subgroups all observed. This heterogeneity at the single-cell level means that the net direction of the population-level effect cannot be predicted from any one recording configuration; it can only be resolved by simultaneously recording from a spatially distributed sample of the network. To our knowledge, no published study has characterized DOI exposure effects on firing, bursts, and functional connectivity at the network level in cortical cultures with a within-culture ketanserin arm — a 5-HT2AR antagonist [24,25] — to probe receptor involvement.

Here we use spatially resolved, multiplexed MEA recordings to characterize post-acute DOI exposure effects on network activity and functional connectivity in dissociated rat cortical cultures, with a within-culture ketanserin arm (Figure 1). We asked three questions. First, how does acute DOI exposure alter spontaneous firing and burst dynamics at the network level, given that DOI drives structural plasticity and excitatory synaptic remodeling in cortical neurons [6,12] but single-cell recordings yield contradictory predictions [22,23]? Second, does acute DOI exposure alter functional connectivity network structure — the pattern of pairwise temporal correlations, not synaptic wiring — beyond what can be explained by firing-rate changes alone? If DOI shifts these functional connectivity patterns, as the REBUS framework’s prediction of decreased modular differentiation under psychedelics [26] would suggest, then graph-theoretic metrics should shift in a direction not fully reducible to per-channel rate. Third, how does ketanserin incubation alter the DOI network response? We interpret this arm as an antagonist exposure condition consistent with 5-HT2AR involvement, not as a receptor-specific mediation test.

**Figure 1.**
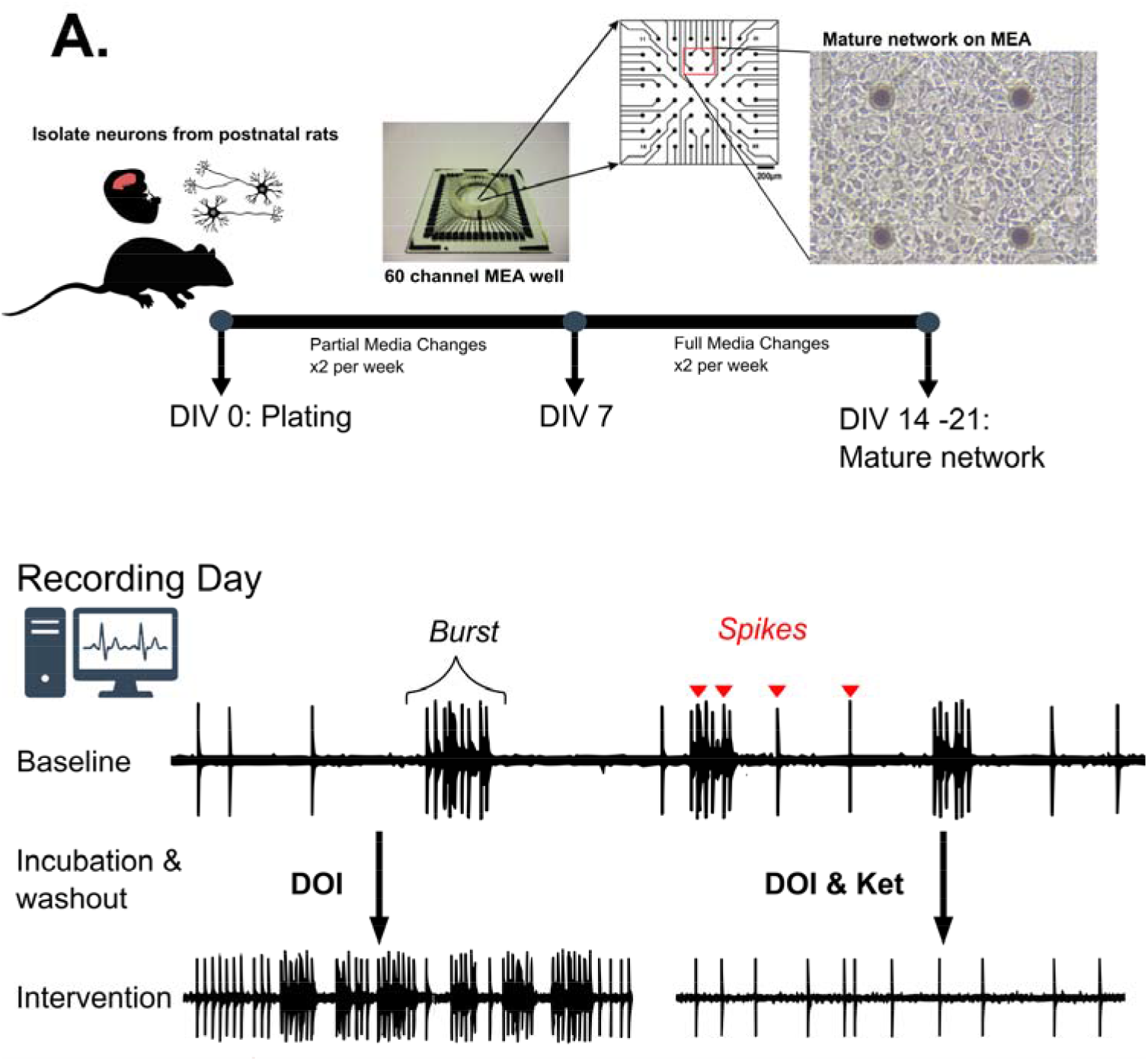
Experimental design. Top: primary cortical neurons were isolated from postnatal rat pups and plated onto microelectrode arrays (MEAs). Cultures were maintained until a mature spontaneously active network had formed (DIV 14-21). Insets show the MEA chip, the 8 × 8 electrode grid layout, and a phase-contrast micrograph of neurons on the electrode array. Bottom: on the recording day, each well contributed a within-culture repeated-measures design. A baseline recording of spontaneous activity was acquired, followed by pharmacological intervention, washout, and an experimental recording of equal duration. Two pharmacological arms were tested: DOI alone (n = 6 wells) and DOI with ketanserin incubation (n = 3 wells). Representative filtered voltage traces from a single channel illustrate baseline activity. After acute exposure, the DOI arm shows increased burst activity, whereas the DOI + ketanserin arm shows suppressed bursting.

## Materials and Methods

All animal-derived primary tissue used in this study was obtained from collaborators within the University of Wisconsin-Madison and used in accordance with institutional guidelines and approved animal-use protocols. All analyses were performed using custom MATLAB scripts (R2023B, MathWorks, Natick, MA) built on the TDT MATLAB SDK for data import and the Signal Processing Toolbox.

### Primary cortical neuron culture

Primary rat cortical neurons were plated onto MEAs following established cortical MEA protocols [14,15,16]. Dissociated cortical preparations of this type are typically composed of glutamatergic pyramidal neurons (∼80–90 %), GABAergic interneurons (∼10–20 %), and glia, with the GABAergic fraction reaching adult-like proportions within the first two weeks in vitro [27,28]. These proportions describe typical cortical preparations and microcircuit composition rather than measurements made directly in the present wells. We use the term *well* to refer to a single MEA chip and its dissociated cortical culture, which is the experimental unit for all paired analyses. Briefly, MEAs were cleaned with 1 % Tergazyme (3-24 h soak), rinsed thoroughly with reagent-grade water, sterilized with 70 % ethanol and ultraviolet light, and incubated overnight at 37 °C / 5 % CO□ under a 50 µL droplet of phosphate-buffered saline (PBS) over the electrode region. Substrate was prepared as 0.1 mg/mL poly-D-lysine (PDL) and 4 µg/mL laminin (Gibco, catalog #23017-015). Substrate droplets (50 µL) were applied to the electrode region, incubated at 37 °C / 5 % CO □ for 60 min, and washed three times with PBS, leaving the final wash on until plating.

Dissociated cells were received at approximately 10 × 10□ cells/mL, manually counted, and diluted to approximately 4 × 10□ cells/mL in unwarmed plating medium. After aspirating the PBS wash, a 50 µL drop of the diluted suspension was placed on each MEA within the established droplet area, yielding approximately 8,333 cells per MEA. Cultures were incubated for 4-24 h before the plating medium was replaced with at least 1 mL of warmed maintenance medium per MEA. Plating medium consisted of Neurobasal Plus supplemented with 1× GlutaMAX and 10 % fetal bovine serum; maintenance medium consisted of Neurobasal Plus supplemented with 1× B27 Plus and 1× penicillin/streptomycin. Half-volume media changes were performed twice weekly during the first week, followed by full-volume changes thereafter. Cultures were maintained at 37 °C / 5 % CO□ and first showed reliable spontaneous activity at approximately DIV 7, consistent with established timelines for this preparation.

### Multi-electrode array recording

Recordings were performed on commercial passive MEA chips from Multi Channel Systems (MCS) (Reutlingen, Germany), product codes 60MEA100/10iR-ITO-GR and 60MEA100/10iR-Ti-GR. Both variants share the same 60-position 8 × 8 grid with the four corners absent and one internal reference electrode, yielding 59 recording electrodes, an inter-electrode pitch of 100 µm, and an electrode diameter of 10 µm; they differ only in the contact track metal (indium tin oxide vs. titanium). Both variants were used across recording sessions and data from the two were pooled for all subsequent analyses.

Signals were acquired through a Tucker-Davis Technologies (TDT, Alachua, FL) TDT MZ60 MEA headstage amplifier streamed through PZ5 neurodigitizer amplifier and RZ5P base processor at the native TDT sampling rate (24.4 kHz). The TDT data store was configured to record up to 64 channels; the 60 MCS MEA positions map to PZ5 channels [1:15, 17:31, 33:47, 49:63], with PZ5 channels 16, 32, 48, and 64 grounded. The internal reference electrode (MCS/PZ5 channel 15) was excluded before rate and connectivity analyses, leaving 59 recording electrodes. Channel-to-grid mapping was verified against the MCS standard 60-electrode layout document (Supplementary Figure S1; SM1). Recordings were carried out at 37 °C inside an enclosed chamber to minimize temperature drift and contamination.

### Pharmacological intervention

Each experimental well contributed a within-well repeated-measures recording: an approximately 10-minute baseline of spontaneous activity, followed by pharmacological intervention, washout, and an approximately equal-duration experimental recording. Two experimental arms were collected: 1. DOI arm (n = 6 wells). DOI was applied at a final bath concentration of 10 µM for ∼20 min, followed by washout and experimental recording. 2. DOI + Ketanserin arm (n = 3 wells). Wells were co-incubated with ketanserin and DOI, each at 10 µM, for 10 min before washout and experimental recording.

### Signal processing

Raw broadband signals were loaded using the TDT MATLAB SDK and processed through the following filter chain:

1. Power-line notch stack. Cascaded zero-phase IIR notch filters at 60 Hz and all integer harmonics up to 780 Hz, with quality factor Q = 35, were applied to suppress line-noise structure before spike-band filtering; zero-phase filtering was used to preserve waveform symmetry. 2. Spike-band bandpass. A 4th-order Butterworth bandpass with cut-off frequencies 300 Hz and 2,500 Hz, applied with zero-phase filtering.

### Spike detection

Per-channel spike detection used the TDT MATLAB SDK threshold detector (TDTthresh) with the following parameters: per-channel noise estimation, negative-going polarity, a threshold factor of 5.0 standard deviations (SD), and a 1 ms post-detection refractory window [29,30]. Detected spike timestamps were sorted and cached for downstream analysis.

### Burst detection

Bursts were extracted from each channel’s sorted spike train using an interspike-interval (ISI) state machine [14,29,31,32]. Parameters were: within-burst ISI threshold = 100 ms (an ISI <= 100 ms initiates or continues a burst) [14,33], termination ISI threshold = 200 ms (an ISI > 200 ms ends a burst), and minimum number of spikes per burst = 5 [33]. Bursts that ran to the end of the recording were closed at the last spike if they met the minimum-spike rule.

### Rate metrics

Following Brofiga et al. [33], for each electrode of each recording we computed mean firing rate (MFR; spikes per minute) and mean burst rate (MBR; bursts per minute), each defined as the event count divided by the recording duration in minutes. To complement MBR with a burst-timing measure, inter-burst interval (IBI; s) was computed as the median interval between consecutive burst onsets on each electrode. Electrodes with MFR < 0.1 spikes/s (6 spikes/min) were treated as inactive for MFR summaries [18,33]. MBR was summarized among electrodes that were already bursting at baseline (baseline MBR > 0), while the number of active electrodes and the number of bursting electrodes were retained as descriptive participation summaries. IBI summaries included electrodes with finite positive values in both baseline and treatment. For each included paired electrode the percent change was computed as:

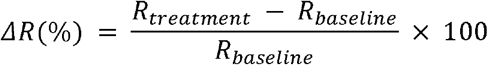

Percent-change summaries used baseline-active electrodes for MFR, baseline-bursting electrodes for MBR, and electrodes with finite positive IBI values in both conditions for the burst-timing metric. Electrodes with no baseline activity for a given metric and non-zero treatment activity were not converted into arbitrarily large percent-change values for the primary intensity summaries; they were instead counted in the corresponding gained-participation category for descriptive visualization.

### Functional connectivity

Pairwise functional connectivity between electrodes was estimated from z-scored, 1 ms-binned spike-count vectors. For each electrode pair (i, j), the Pearson cross-correlation was computed across a lag window of +/-100 ms, and the peak value was taken as the connection weight:

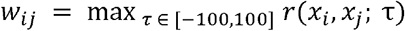

where x□ and x□ are the z-scored binned spike-count vectors for electrodes i and j and tau is the time lag in milliseconds. The lag at which the peak occurred was retained as the peak-lag estimate. The result for each recording is a 59 × 59 symmetric adjacency matrix and a corresponding peak-lag matrix over the recording-electrode set. Any channel whose mean cross-correlation fell below one-tenth of the per-recording median was masked for adjacency-matrix visualization; graph metrics were computed after proportional thresholding at 10 % edge density, so only the strongest pairwise connections entered the binarized graph. The peak cross-correlation approach follows Garofalo et al. (2009) [34] and Poli, Pastore and Massobrio (2015) [35].

#### Spike-time tiling coefficient (STTC)

Because the Pearson cross-correlation on binned spike trains is sensitive to absolute firing rate when rates change sharply between conditions, every pairwise edge was also estimated with the spike-time tiling coefficient (STTC) of Cutts and Eglen (2014) [36], an estimator that is intrinsically rate-corrected. For two spike trains A and B and a synchrony window Δt, STTC is defined as:

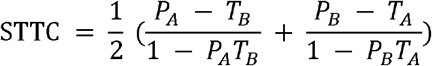

where T for train A is the proportion of the recording duration falling inside ±Δt tiles around any spike in train A; T for train B is the analogous quantity for B; P for train A is the proportion of spikes in A that fall within ±Δt of any spike in B; and P for train B is the analogous quantity for B. STTC is bounded on [-1, 1] and equals zero in expectation for two independent Poisson trains regardless of their rates, which is the property that makes it rate-corrected. We used a synchrony half-window of Δt = 50 ms, matching the Cutts and Eglen implementation used here [36]. STTC adjacency matrices were computed for every recording and binarized at the same 10 % edge density used for the Pearson analysis, allowing direct estimator-by-estimator comparison of every graph metric.

#### Network-level metrics

Graph-theoretic summaries were computed after binarizing each adjacency matrix at a target edge density of 10 % using proportional thresholding [37,38]. Metrics include edge density, clustering coefficient [39] (averaged across nodes), characteristic path length (on the largest connected component), global efficiency [40], modularity Q [41] (greedy agglomerative community detection), and small-worldness σ [42]. Characteristic path length was defined as:

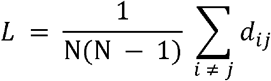

where d□□ is the shortest binary-graph path between electrodes i and j and N is the number of graph nodes in the largest connected component. Modularity was defined as:

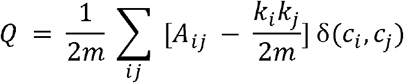

where AD□ is the binary adjacency entry, k□ and k□ are node degrees, m is the total number of edges, c□ and c□ are community assignments, and δ is 1 when two nodes are assigned to the same community and 0 otherwise. Small-worldness was quantified as:

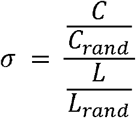

where C and L are the observed clustering coefficient and characteristic path length, and the corresponding random-null C and L values are averaged over 50 Erdos-Renyi graphs matched on node and edge count. All metric definitions follow Rubinov and Sporns (2010) [37,43].

### Topographical mapping

Per-electrode firing-rate, burst-rate, and connectivity-derived metrics were projected onto the standard MCS 60-electrode 8 × 8 layout for visual inspection. Channel-to-grid mapping was verified against the MCS standard 60-electrode layout document (Supplementary Figure S1).

### Statistical analysis

The analysis is structured around three research questions, matching the Introduction: Q1 asks how DOI exposure alters firing and burst dynamics; Q2 asks how acute DOI exposure alters functional connectivity network structure and whether that graph-metric shift is separable from firing-rate change; Q3 asks how incubation with ketanserin alters the DOI network response.

#### Q1

DOI effects on firing and burst dynamics (within-culture repeated-measures, n = 6 wells). The primary rate endpoints were MFR, MBR, and IBI. Primary DOI-arm inference used exact Wilcoxon signed-rank tests on per-well medians, with Benjamini-Hochberg (BH) adjustment across the prespecified rate endpoints [44]. We also report the number of wells changing in the same direction as a descriptive concordance summary, together with paired Hedges’ g using an average-SD denominator (hereafter Hedges’ g) [45]. The two-level hierarchical bootstrap (10,000 iterations, seed = 1) [46] is reported in SM5. Bootstrap p-values remain Supplementary Information sensitivity summaries and are not used for primary significance calls.

#### Q2

DOI effects on functional connectivity network structure (within-culture repeated-measures, n = 6 wells). The primary functional-network endpoints were characteristic path length, modularity Q, and small-worldness σ computed from Pearson cross-correlation adjacency matrices thresholded at 10 % edge density. The same exact well-level Wilcoxon framework is used for these metrics, with BH-adjusted *p* reported after BH correction across those endpoints. The principal rate-control analysis is the spike-time tiling coefficient (STTC) [36], which is intrinsically rate-corrected. The three functional-network metrics are recomputed using STTC-derived adjacency matrices in parallel with the Pearson cross-correlation analysis. Convergent direction across the two estimators is taken as evidence that the graph-metric change is not simply an artifact of the rate sensitivity of Pearson.

#### Q3

Ketanserin + DOI exposure arm (n = 3 wells). The ketanserin arm is reported descriptively using directional concordance, per-well median changes, Hedges’ g, and hierarchical-bootstrap confidence intervals. No formal inferential or between-arm hypothesis test was used for the ketanserin arm. Interpretation is framed as a pharmacological exposure result consistent with, but not proof of, 5-HT2AR involvement.

Sample-size justification. Sample sizes follow established precedent in MEA pharmacology and exploratory cortical-slice MEA studies, where within-preparation designs often use small numbers of independent preparations or wells [19,30,38]. Our DOI arm (n = 6 wells, >200 active electrodes) was analyzed with exact well-level tests as the primary inference and a supplementary hierarchical bootstrap to summarize uncertainty in the nested electrode data [46]. The ketanserin arm (n = 3) is reported descriptively with effect sizes and confidence intervals rather than treated as a powered confirmatory test.

All random seeds are fixed (seed = 1) for reproducibility.

## Results

### DOI increases firing and accelerates burst timing

Across the six DOI recordings, baseline activity displayed the characteristic dissociated-cortex signature of tonic multi-unit firing punctuated by short population bursts [14,15,16]. Because synchronized burst events are known to contribute strongly to pairwise temporal correlations in dissociated cortical cultures [14,47], MFR, MBR, and IBI features were quantified alongside functional connectivity [33]. The median baseline MFR among active electrodes was 73.7 spikes/min, and the median baseline MBR among baseline-bursting electrodes was 5.44 bursts/min. Following acute DOI exposure, MFR increased across all six wells (median within-well change = +41.6 %, Hedges’ g = 0.42, bootstrap 95 % CI [3.6 %, 144.9 %], 6/6 wells increased, BH-adjusted *p* = 0.047; Figure 2c). MBR among baseline-bursting electrodes increased directionally (median within-well change = +103.5 %, Hedges’ g = 0.72, bootstrap 95 % CI [2.1 %, 125.4 %], 5/6 wells increased, BH-adjusted *p* = 0.094; Figure 2d), and the IBI shortened across all six wells (median within-well change = -48.1 %, Hedges’ g = -1.35, 6/6 wells shortened, BH-adjusted *p* = 0.047; Figure 2e). Descriptive participation summaries showed near-ceiling active-electrode counts at baseline and after DOI exposure, whereas bursting-electrode counts varied by well (Figure 2f-h). Thus DOI exposure primarily increased firing and accelerated burst timing among participating electrodes rather than recruiting additional active electrodes.

**Figure 2.**
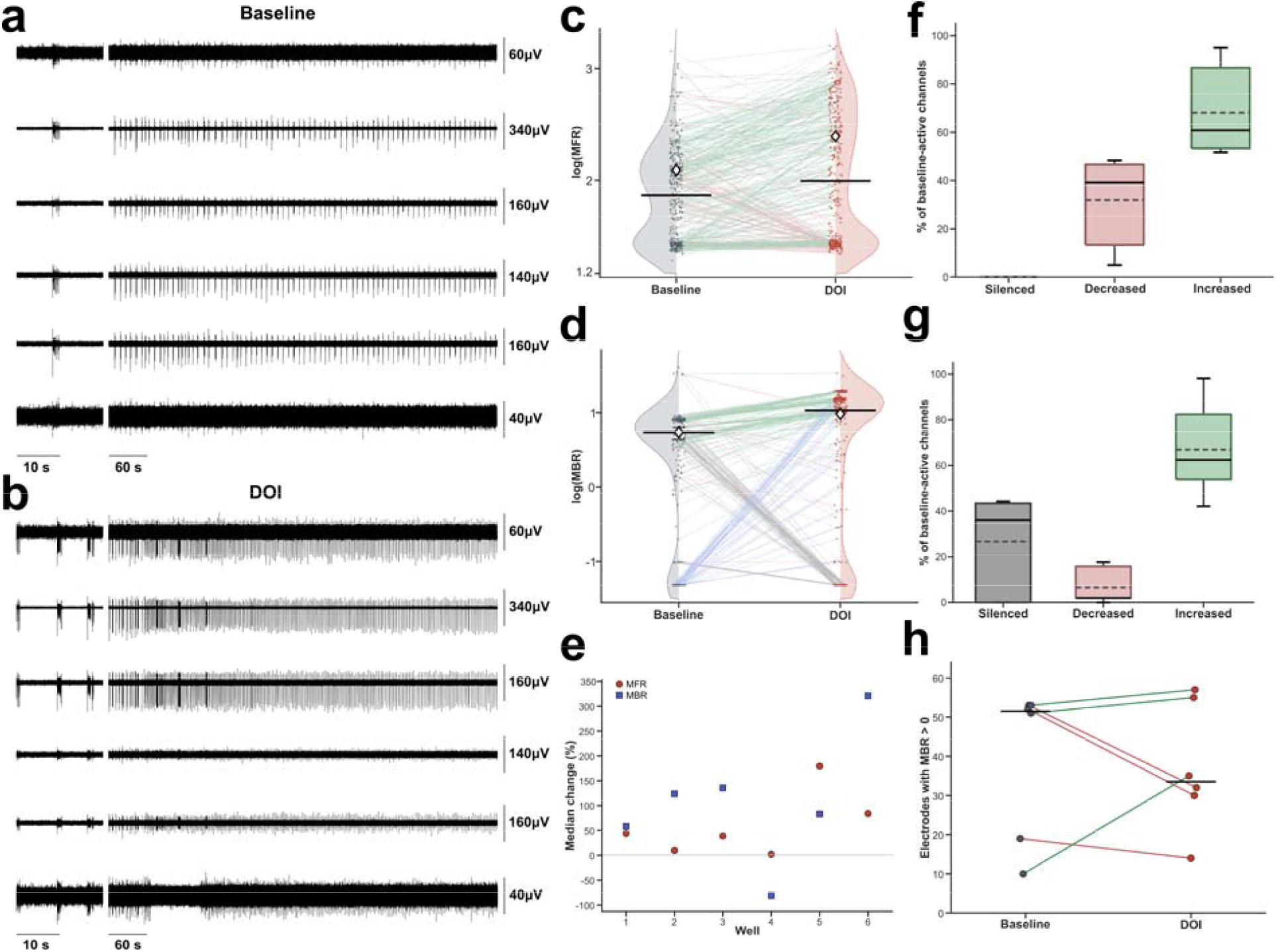
Acute DOI exposure increases firing and accelerates burst timing in cortical cultures. (a) Baseline filtered voltage traces from representative electrodes of one within-culture repeated-measures DOI recording, shown at 10 s and 60 s time scales with per-electrode amplitude scale bars. (b) Post-acute DOI traces from the same recording and electrodes, using the same time scales and amplitude scaling convention. (c) Paired baseline-DOI MFR distributions for active electrodes (MFR >= 0.1 spikes/s), displayed as log-transformed values following Brofiga et al.; shaded half-density envelopes show condition-wise electrode distributions, dots mark individual electrodes, paired lines connect matched electrodes, solid black ticks mark pooled medians, and open diamonds mark pooled means. Paired-line colors encode categorical channel changes: green = increased, red = decreased, blue = newly active or newly bursting, and grey = unchanged or silenced. (d) Paired MBR distributions for electrodes with detectable bursting in either condition, displayed as log-transformed values with zero MBR values placed at a small plotting floor for visualization only. (e) Per-well median percent change for MFR, MBR, and IBI. (f) Active-channel state summary, expressed as a percentage of baseline-active channels. (g) Baseline-active burst-channel state summary. (h) Paired count of electrodes with MBR > 0 at baseline and after DOI exposure. Statistics use raw values; inference is led by exact Wilcoxon tests on per-well median percent change for MFR, MBR, and IBI, with participation panels shown descriptively.

### DOI shifts functional network structure toward integration

Network-level graph metrics computed from Pearson cross-correlation adjacency matrices thresholded at 10 % edge density [37,38] suggested an integration-oriented shift in functional-network structure following acute DOI exposure (Figure 3). The three functional-network metrics were characteristic path length, modularity Q, and small-worldness σ. Characteristic path length shortened in 5/6 wells (median change = -0.17, Hedges’ g = -0.93, bootstrap 95 % CI [-.68, -0.07], BH-adjusted *p* = 0.031; Figure 3d1). Modularity Q decreased in 5/6 wells (median change = -0.05, Hedges’ g = - 0.40, bootstrap 95 % CI [-0.18, -0.004], BH-adjusted *p* = 0.031; Figure 3d2), and small-worldness σ increased in 5/6 wells (median change = +0.81, Hedges’ g = +1.18, bootstrap 95 % CI [0.09, 2.27], BH-adjusted *p* = 0.031; Figure 3d3). Thus all three metrics moved in the predicted direction and remained significant after BH correction, with the standardized effect-size summary emphasizing the largest magnitudes for characteristic path length and small-worldness (Figure 3e). Spatial-decay plots show how the strongest surviving correlations are distributed across inter-electrode distance, providing a topology check complementary to the graph metrics (Figure 3f).

**Figure 3.**
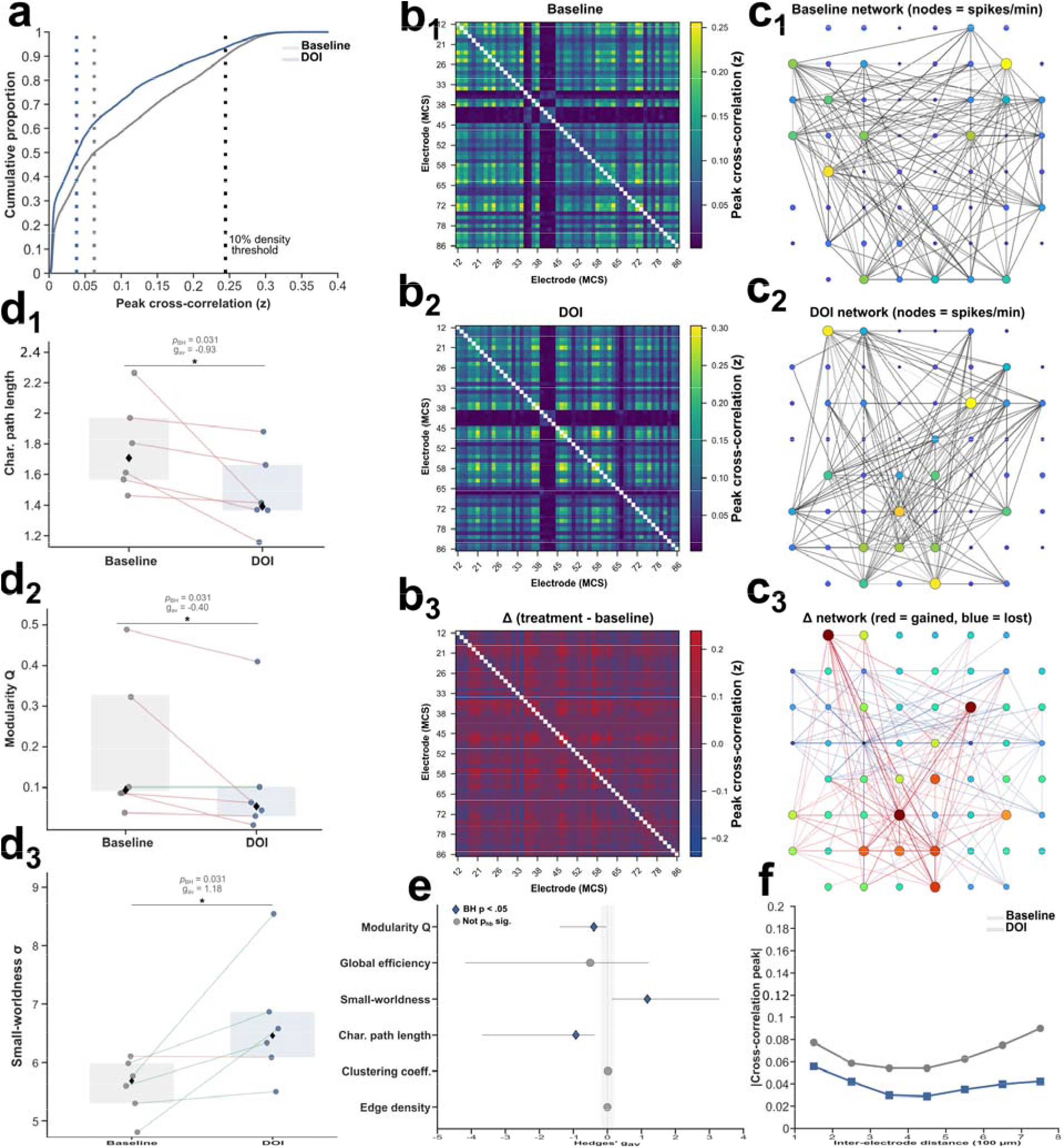
Acute DOI exposure shifts functional network structure. (a) Cumulative distribution of all pairwise Pearson cross-correlation peak values pooled across six DOI recordings. The DOI curve (blue) shifts relative to baseline (grey); because graph metrics are computed after proportional thresholding at 10 % edge density, the graph-metric analysis captures how the strongest connections are redistributed rather than a simple change in total edge count. Dotted vertical lines mark the condition-specific 10 % edge-density thresholds used for graph binarization. (b1-b3) Pairwise Pearson cross-correlation matrices for a single exemplar recording, sorted by MCS electrode label: baseline (b1), post-acute DOI (b2), and post-exposure-minus-baseline difference (b3; red = strengthened, blue = weakened). (c1-c3) Functional network graphs drawn on the 8 × 8 electrode layout for baseline (c1), post-acute DOI (c2), and the edge-change network (c3). Node size and color encode per-electrode firing rate (spikes/min); baseline and post-acute DOI edges represent the top 10 % of the Pearson cross-correlation distribution, and the difference network shows gained (red) and lost (blue) edges. (d1-d3) Paired well-level graph metrics for characteristic path length (d1), modularity Q (d2), and small-worldness σ (d3). Each line connects one within-culture baseline-DOI recording; diamonds mark group medians and shaded bars show IQRs. Characteristic path length and modularity Q decreased in 5/6 wells, whereas small-worldness increased in 5/6 wells (BH-adjusted p = 0.031 for each prespecified metric). (e) Standardized effect-size summary for functional-network metrics. Points show Hedges’ g, horizontal lines show bootstrap 95 % CIs, and marker shape indicates whether the exact well-level Wilcoxon test was BH-significant. (f) Spatial decay of functional connectivity, showing mean |cross-correlation peak| as a function of inter-electrode distance (100 µm units), for baseline and post-acute DOI conditions. Complete metric tables and bootstrap sensitivity summaries are reported in SM4-SM5, density sweeps in SM3, and STTC rate-control analyses in SM2.

### Ketanserin suppresses burst dynamics

In the three DOI + ketanserin wells, the central observation was that acute ketanserin + DOI exposure reduced MBR and prolonged burst timing while leaving per-channel MFR comparatively modest and inconsistent. Per-well MFR showed a small and inconsistent shift across the three wells (median within-well change = -4.3 %, Hedges’ g = -0.88, bootstrap 95 % CI [-25.0 %, 3.3 %], 1/3 wells increased; Figure 4c). MBR among baseline-bursting electrodes decreased in 3/3 wells (median within-well change = -90.5 %, pooled paired-electrode Hedges’ g = -0.07, bootstrap 95 % CI [-100.0 %, -83.2 %], Figure 4d), and the IBI increased markedly among electrodes with measurable burst timing in both conditions (median within-well change = +326.8 %, Hedges’ g = +1.74, 3/3 wells increased; Figure 4e). Descriptive participation summaries showed preserved active-electrode participation and reduced bursting-electrode participation (Figure 4f-h). The population-level MBR summary therefore describes the median of a heterogeneous electrode-level response, not a uniform suppression.

**Figure 4.**
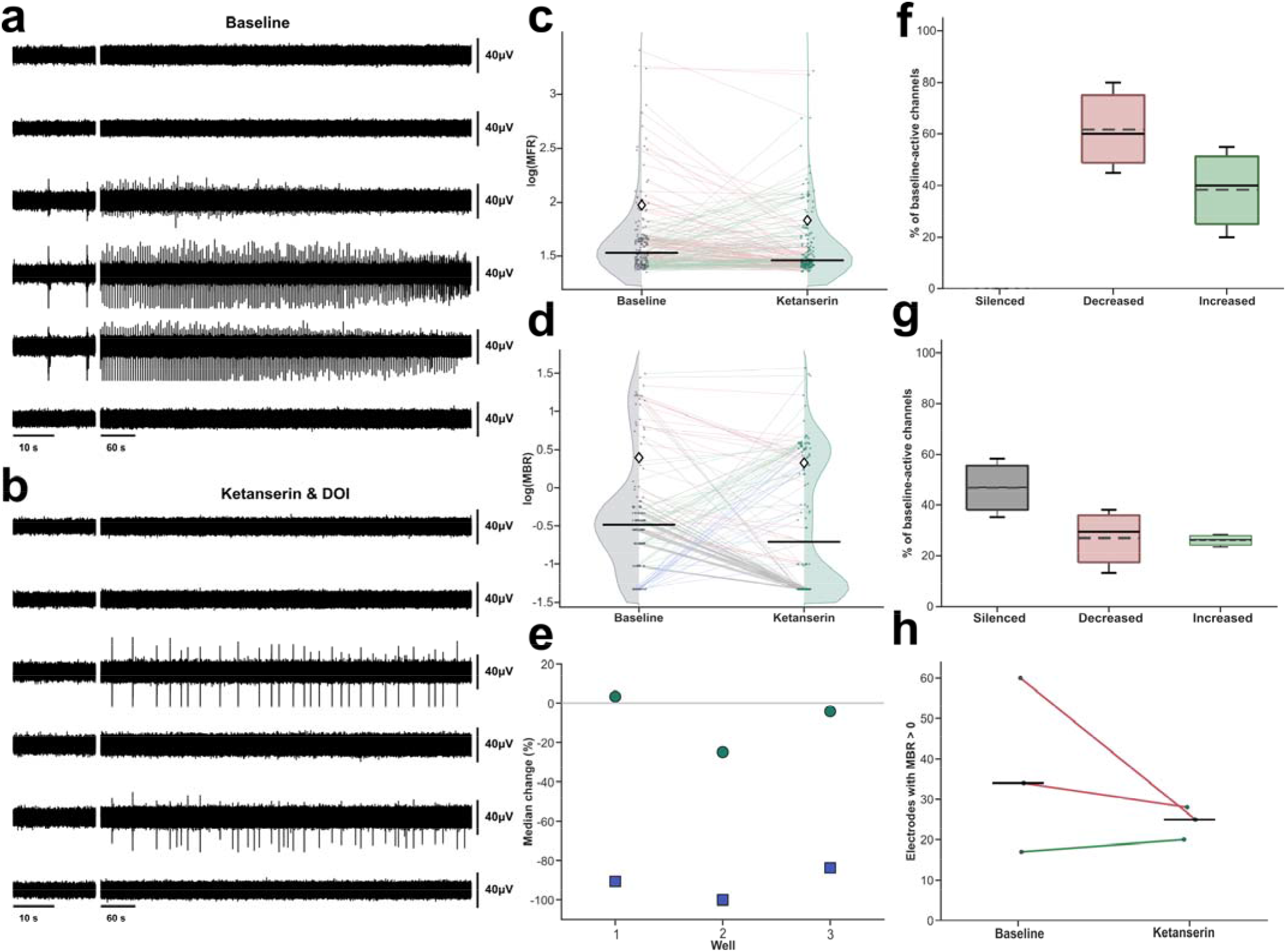
Ketanserin + DOI exposure reduces MBR and prolongs IBI while preserving active-electrode participation. (a) Baseline filtered voltage traces from representative electrodes of one DOI + ketanserin recording, shown at 10 s and 60 s time scales with per-electrode amplitude scale bars. (b) Post-acute ketanserin + DOI traces from the same recording and electrodes, using the same time scales and amplitude scaling convention. (c) Paired baseline-post-exposure MFR distributions for active electrodes (MFR >= 0.1 spikes/s), displayed as log-transformed values following Brofiga et al. Shaded half-density envelopes, dots, paired lines, solid median ticks, and open mean diamonds follow Figure 2; paired-line colors encode categorical channel changes rather than intensity-scaled effect magnitude. (d) Paired MBR distributions for electrodes with detectable bursting in either condition, displayed as log-transformed values with zero MBR values placed at a small plotting floor for visualization only. (e) Per-well median percent change for MFR, MBR, and IBI. (f) Active-channel state summary, expressed as a percentage of baseline-active channels. (g) Baseline-active burst-channel state summary. (h) Paired count of electrodes with MBR > 0 at baseline and after ketanserin + DOI exposure. Statistics use raw values. The ketanserin arm is reported descriptively because n = 3 wells.

**Table 1.** Summary of firing and burst metrics.

| Arm | Metric | Unit | n wells | n elec. | Baseline median | Treatment median | Median change (%) | BH-adjusted p |
| --- | --- | --- | --- | --- | --- | --- | --- | --- |
| DOI | MFR | spikes/min | 6 | 360 | 73.70 | 98.67 | 41.59 | 0.047 |
| DOI | MBR | bursts/min | 6 | 280 | 5.442 | 10.73 | 103.5 | 0.094 |
| DOI | IBI | s | 6 | 179 | 8.092 | 4.352 | -48.06 | 0.047 |
| Ketanserin | MFR | spikes/min | 3 | 180 | 33.95 | 28.81 | -4.256 |  |
| Ketanserin | MBR | bursts/min | 3 | 128 | 0.327 | 0.197 | -90.54 |  |
| Ketanserin | IBI | s | 3 | 45 | 3.576 | 19.75 | 326.8 |  |

*Note.* Descriptive values are pooled electrode-level summaries using the same inclusion rules as Figures 2 and 4. DOI p values are Benjamini-Hochberg-adjusted exact Wilcoxon signed-rank, computed on per-well median percent change. The ketanserin arm is reported descriptively because $n = 3$ wells.

### Ketanserin produces path-length shortening with estimator-sensitive modularity

The three DOI + ketanserin recordings showed a descriptive graph-metric pattern that partially overlapped with the DOI arm (Figure 5). Given the small sample size and possible off-target ketanserin effects during exposure, this arm is not treated as a powered receptor-specific functional-network test. Edge density was held constant at 10 %. All three wells showed a Pearson path-length decrease (median change = -0.24, Hedges’ g = -0.53, bootstrap 95 % CI [-0.43, -0.18]; Figure 5d1). Modularity Q increased descriptively (median change = +0.04, Hedges’ g = +0.20, bootstrap 95 % CI [-0.17, 0.28]; Figure 5d2), while small-worldness σ rose in all three wells (median change = +1.96, Hedges’ g = +0.65, bootstrap 95 % CI [0.77, 2.27]; Figure 5d3). The standardized effect-size summary shows the same descriptive pattern across the plotted functional-network metrics (Figure 5e).

**Figure 5.**
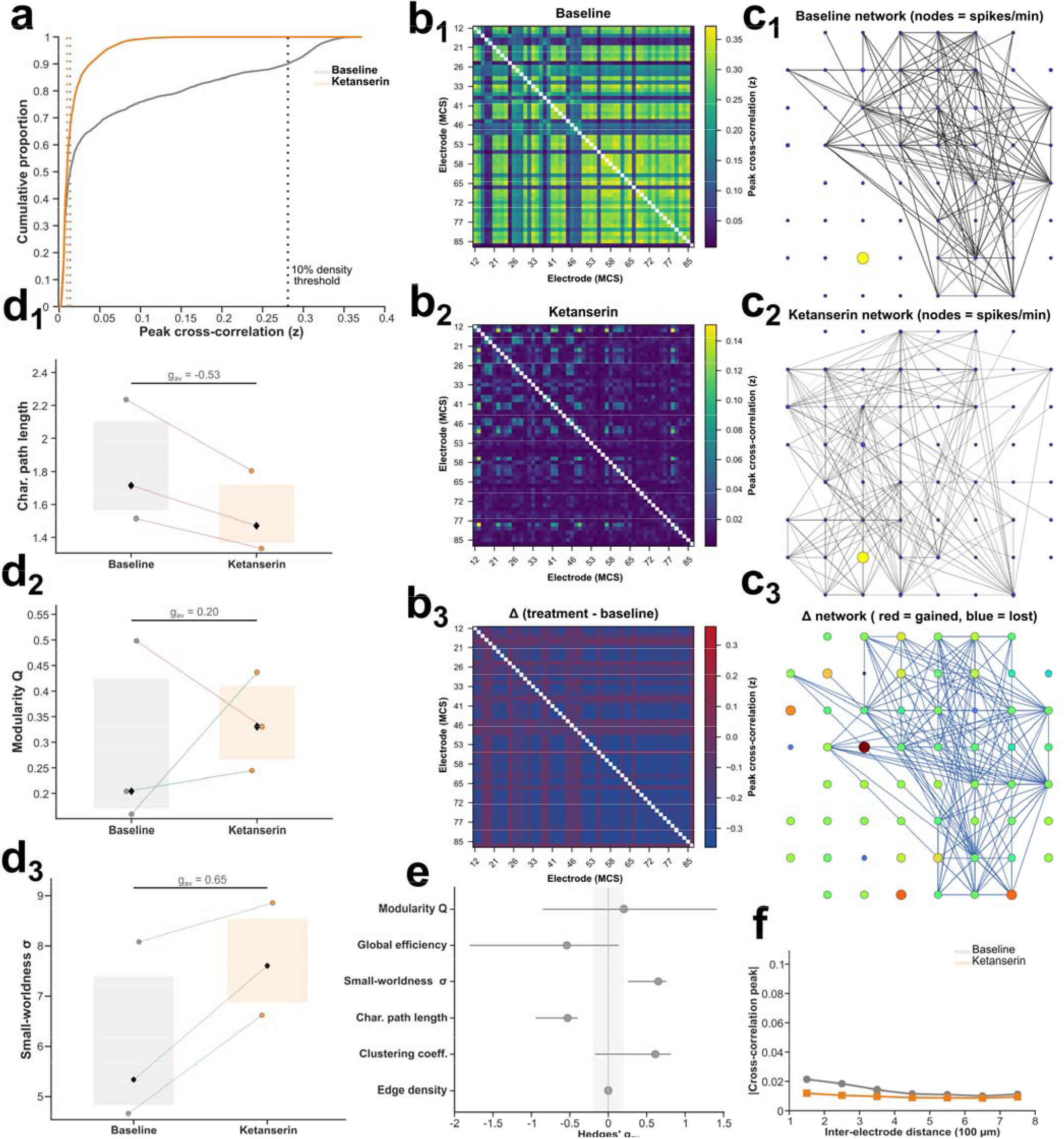
The ketanserin arm shows descriptive shifts in functional-network metrics at n = 3. (a) Cumulative distribution of all pairwise Pearson cross-correlation peak values pooled across three DOI + ketanserin recordings, same format as Figure 3a. The post-acute ketanserin + DOI curve (orange) shifts leftward, consistent with broad reduction of pairwise coupling after the burst-rate collapse; dotted vertical lines mark the condition-specific 10 % edge-density thresholds. (b1-b3) Pairwise Pearson cross-correlation matrices for a single exemplar recording: baseline (b1), post-acute ketanserin + DOI (b2), and post-exposure-minus-baseline difference (b3). (c1-c3) Functional network graphs on the 8 × 8 electrode layout for baseline (c1), post-acute ketanserin + DOI (c2), and the edge-change network (c3), same format as Figure 3c1-c3. The ketanserin difference network is dominated by lost edges, consistent with reduced burst-coupled synchrony. (d1-d3) Paired well-level graph metrics for characteristic path length (d1), modularity Q (d2), and small-worldness σ (d3). All three wells showed a path-length decrease (Pearson medians 1.71 to 1.47, range of decreases 0.18-0.43), modularity Q increased descriptively, and small-worldness increased in all three wells. (e) Standardized effect-size summary for functional-network metrics, same format as Figure 3e. (f) Spatial decay of functional connectivity, baseline vs post-acute ketanserin + DOI conditions.

### Functional-network sensitivity analyses and estimator checks

Complementary functional-network metrics, bootstrap sensitivity estimates, density sweeps, and estimator comparisons are collected in the Supplementary Information (SM2-SM5; Supplementary Tables S1-S6). The principal rate-control check recomputed the graph metrics with STTC, a spike-timing estimator designed to reduce firing-rate bias [36]. In the DOI arm, path-length shortening remained directionally negative under STTC, although the effect was weaker than the Pearson estimate and present in 4/6 wells rather than 5/6 wells (Pearson median Δ = -0.170; STTC median Δ = -0.100; SM2 and Supplementary Table S1). In the ketanserin arm, path-length shortening was preserved under STTC in all 3 wells (STTC medians 1.88 to 1.37), while the modularity Q signal reversed sign between estimators (Pearson Δ = +0.040; STTC Δ = -0.069; SM2 and Supplementary Table S2). These estimator comparisons identify path length as the most stable functional-network axis and modularity, especially in the ketanserin arm, as estimator-sensitive. Together these analyses suggest that the functional-network results carry information about network state that is not fully reducible to per-electrode MFR; the strength of this conclusion is bounded by estimator sensitivity and sample size.

## Discussion

The clearest DOI effect was a shift toward a more active, faster-bursting network state. DOI increased spontaneous MFR (median within-well change = +42 %, 6/6 wells positive, BH-adjusted *p* = 0.047), shortened IBI (median within-well change = -48 %, 6/6 wells negative, BH-adjusted *p* = 0.047), and directionally increased MBR among baseline-bursting electrodes (median within-well change = +104 %, 5/6 wells positive, BH-adjusted *p* = 0.094). DOI functional-network metrics showed a convergent but sensitivity-limited pattern toward shorter characteristic path length, lower modularity Q, and higher small-worldness σ, consistent with a possible shift in the integration-segregation balance [37,48]. Ketanserin incubation with DOI reduced MBR, prolonged IBI, and produced a descriptive path-length shortening that was directionally consistent across all 3 wells. Path-length shortening was preserved in both arms under the rate-corrected STTC estimator, arguing against a change simply identical to per-channel firing rates while remaining bounded by sample size and estimator sensitivity.

### Spatially resolved functional characterization of DOI in cortical networks

Prior in-vitro DOI work operated at the single-cell level: Hu et al. (2016) [22] showed via patch clamp that DOI *depresses* spontaneous firing of individual cultured cortical neurons through 5-HT2A/2C activation, while in the same preparation DOI nearly doubled mEPSC frequency without changing mEPSC amplitude (3.9 → 7.4 Hz) — a synaptic effect that would, by excitation-inhibition balance alone, favor enhanced spontaneous firing. The structural-plasticity literature has separately documented dendritic growth and spinogenesis on single neurons [6,7,12]. The current experimental design should be interpreted as a post-acute network state initiated by DOI exposure rather than direct receptor occupancy during recording. This follows the work by Ly et al. demonstrating that psychoplastogen stimulation and later growth can be temporally dissociated after ligand removal, while DOI-specific work shows that DOI can engage cortical plasticity and Trk-related trophic-signaling pathways [6,7,12]. Our MEA recordings provide direct evidence at the network level: MFR increases by +42 %, MBR shifts upward directionally by +104 %, IBI shortens by -48 %, and the functional network shifts toward shorter characteristic path lengths.

The difference between acute single-cell depression (Hu et al.) and post-acute network-level excitation in the present study is not, however, necessarily contradictory. Puig et al. (2003) [23] showed in the published article that systemic DOI produces heterogeneous effects across 56 identified mPFC pyramidal neurons, including excitatory (38 %), inhibitory (30 %), and unaffected (32 %) subgroups. Our MEA recordings capture the aggregate of these opposing single-cell effects: in a recurrently connected dissociated network, the excitatory component may dominate the post-exposure population state, yielding a net increase in MFR and accelerated burst timing despite mixed electrode-level responses. Whether these subpopulations correspond to distinct cell types as the preparation contains a mix of glutamatergic pyramidal cells, GABAergic interneurons, and glia, with 5-HT2A expression predominantly on pyramidal neurons [23] or to variation in receptor expression density across the cultured network remains to be determined. This distinction between single-cell heterogeneity and population-level outcome could only be made by simultaneously recording from a spatially distributed sample of the network, which is the specific gap our preparation fills.

### Ketanserin and 5-HT2AR involvement

The ketanserin result merits specific interpretation. Ketanserin is a standard antagonist used to attenuate DOI- and 5-HT-linked responses in neuronal preparations [22,49,50]. At the 10 µM bath concentration used during exposure, however, ketanserin should be interpreted as a saturating antagonist condition rather than a receptor-selective dose: it is nearly 3000-fold above ketanserin’s reported 5-HT2A K□(∼3.5 nM) [24], and ketanserin also engages 5-HT2C, α□-adrenergic, H□, and D□ receptors [25]. With that caveat, the combined arm strongly reduced population bursting in 3/3 wells while channel-level firing rate showed only a small and inconsistent shift (per-well median -4.3 %). The dissociation between population-burst reduction and modest channel-level firing-rate change is the central observation: ketanserin incubation with DOI, whether acting through 5-HT2A blockade alone or broader receptor engagement at this dose, was associated with reduced recurrent burst initiation. Because synchronized population bursts strongly shape pairwise temporal correlations in dissociated cortical cultures [14,47], reducing them can reshape functional connectivity network structure — including path length — even without a large change in channel-level firing rate. This pattern is consistent with 5-HT2AR involvement in recurrent excitatory recruitment, but it does not establish receptor-specific mediation.

One speculative mechanism is constitutive 5-HT2A signaling. Ketanserin has documented inverse agonist properties at 5-HT2A in recombinant systems [25], constitutive 5-HT2A activity has been demonstrated behaviorally after near-complete serotonin depletion [51], and recent human prefrontal cortex membrane work shows complex ketanserin-driven G-protein coupling at 5-HT2A [52]. Because the dissociated cortical preparation lacks serotonergic cell bodies — which reside exclusively in the brainstem raphe nuclei — the burst-rate suppression associated with ketanserin exposure could reflect reduced constitutive 5-HT2A signaling rather than competitive displacement of an endogenous agonist. This mechanism remains untested in the present preparation.

### Functional network structure and integration-segregation balance

The three functional-network metrics following acute DOI exposure point toward an altered integration-segregation balance: characteristic path length shortened, modularity Q decreased, and small-worldness σ increased [37,48]. The DOI-associated increase in MBR (+104 % among baseline-bursting electrodes) may create cross-module correlations that bridge formerly segregated communities and shorten the average functional path between electrode pairs, although this mechanistic explanation remains inferential. The convergence of path length, modularity, and small-worldness in the expected direction suggests that DOI exposure altered functional network structure, but the sample size leaves this result underpowered for a strong confirmatory claim. Within these metrics, path length was the most stable estimator-controlled axis, because its direction was preserved under STTC in both pharmacological arms, whereas modularity was more estimator-sensitive in the ketanserin arm. The ketanserin arm produced path-length shortening in a different rate context: by reducing the burst-correlation structure that dominates pairwise coupling in dissociated cultures [14,47], the residual tonic connectivity can yield shorter paths even without an increase in MFR. This dissociation is graph-theoretically plausible because path length is sensitive to shortcut-like edges and to which pairwise connections survive proportional thresholding [37,39]. The convergence of both arms on path-length shortening despite opposite rate effects, together with persistence of the path-length effect under the rate-corrected STTC estimator, supports path length as the most robust functional-network observation in this dataset. It should be interpreted as evidence for a functional-connectivity change not fully reducible to a rate artifact.

This profile aligns with the REBUS model’s prediction that psychedelics flatten the cortical free-energy landscape, enabling neuronal dynamics to escape modular basins of attraction [26]. In line with the REBUS framework, a recent international mega-analysis of acute psychedelic rsfMRI effects in humans provides a useful large-scale counterpart: Girn et al. (2026) found that acute psychedelics consistently shift functional coupling toward greater between-network integration across association, sensory, and subcortical systems [53]. The spatial scale and preparation differ fundamentally from the present MEA recordings, but both datasets point toward increased functional integration. In our reduced cortical preparation, DOI-associated path-length shortening may therefore represent a local circuit analogue of increased cross-network integration. Recent behavioral studies further show that a single dose of DOI produces time-dependent changes in cognitive flexibility, with improvements in reversal learning emerging one week after administration [54], suggesting that the connectivity shift we observe in vitro may represent a circuit-level correlate of the behavioral plasticity these compounds induce.

### Limitations and future directions

Several considerations limit the strength and scope of the present claims. A first constraint concerns pharmacological and mechanistic specificity. Prior in vitro neuronal electrophysiology has used 10 µM bath ketanserin as a blocking concentration in 5-HT- and DOI-linked preparations [49,50], but that precedent supports the operational antagonist exposure condition rather than receptor selectivity. At 10 µM, off-target receptor activity cannot be excluded as a contributor to the ketanserin findings, because ketanserin is expected to engage 5-HT2C, α□-adrenergic, H□, D□, and potentially other signaling pathways at concentrations far above its 5-HT2A affinity [25,52]. The ketanserin arm should therefore be read as a result consistent with, but not proof of, 5-HT2AR involvement in the DOI-associated effects.

A second constraint is statistical rather than mechanistic. The DOI arm (n = 6 wells) and the ketanserin arm (n = 3 wells) are consistent with in vitro MEA pharmacology precedent [18,19] but small in absolute terms. We accordingly use exact Wilcoxon signed-rank tests on per-well medians as the primary inferential layer for the DOI arm, with sign counts reported as descriptive within-well concordance. The two-level hierarchical bootstrap is reported in SM5 as an uncertainty summary for nested electrode data rather than as a primary significance criterion [46].

Experimental controls impose a further boundary on interpretation. A vehicle-only control arm was not included, so any drift attributable to medium exchange, temperature transients, osmotic perturbation, or washout procedure is absorbed into the drug-exposure effects rather than subtracted. A temporal analysis of population firing rate across the full shared recording duration (∼9.5 min) shows that all three conditions (baseline, DOI, DOI + ketanserin) exhibit a brief transient increase in the first ∼30 s followed by stable firing for the remainder of the recording (Supplementary Figure S8), suggesting that the relative rate profiles are not driven by initial perturbation artifacts. Only a single DOI concentration, a single ketanserin concentration, and a single washout protocol were tested. Recordings during DOI exposure, concentration-response series, and delayed post-washout intervals are natural next steps that would help separate residual pharmacology, and emerging plasticity-linked network states.

The connectivity analysis has its own estimator-dependent limitation. The Pearson cross-correlation on z-scored binned spike counts is sensitive to absolute firing rate when rates change sharply between conditions [36]. To address this rate-confound concern without letting the main figures become an estimator audit, the three functional-network metrics were computed in parallel using the rate-corrected spike-time tiling coefficient (STTC) [36], with the Pearson/10 % graphs shown in Figures 3 and 5 and the estimator comparison reported in SM2. Path-length shortening persisted under STTC in both arms, but the DOI effect was weaker under STTC (4/6 wells; median Δ = -0.100) than under Pearson (5/6 wells; median Δ = -0.170), so the estimator comparison bounds the principal graph-metric direction. The ketanserin modularity Q signal reversed sign between Pearson and STTC, indicating that modularity is estimator-sensitive in this small dataset. Density-sweep analyses across 5-20 % target edge density [37] are reported in SM3. Larger independent-well cohorts will be needed to determine whether the directional functional-network pattern is statistically stable.

The dissociated cortical preparation, while offering pharmacological control and reproducibility, loses the regional identity and laminar architecture of intact cortex. The graph metrics we report therefore reflect the emergent functional network structure of a randomly plated neuronal assembly sampled by MEA electrodes, not the circuit-specific effects that might arise in vivo [19]. In vivo validation with high-density electrophysiology — for example, Neuropixels recordings in acute cortical preparations or freely behaving animals — would test whether the integration-shifted functional-connectivity signature we observe in vitro translates to intact circuits with preserved laminar and long-range connectivity.

Computational approaches offer a complementary path forward. In silico models of neuronal network dynamics [55] could generate predictions about how 5-HT2A-mediated changes in synaptic parameters produce the specific graph-metric signatures we observe, and network inference algorithms applied to temporally binned spike trains [56] may improve estimation of effective connectivity beyond pairwise cross-correlation. Machine learning methods applied to MEA spike-train data represent a further opportunity to decode pharmacological states and predict network responses to novel compounds.

## Conclusion

Together, these recordings show that acute DOI exposure reshapes dissociated cortical cultures at the population-network level: firing increased, bursts accelerated, and functional connectivity shifted toward shorter path length despite estimator and sample-size limits. The ketanserin + DOI arm moved the system into a different rate regime by suppressing population bursting, yet path-length shortening persisted, supporting a network-state interpretation while remaining insufficient to establish receptor-specific mediation. These findings place multiplexed cortical network recordings as a tractable bridge between molecular psychoplastogen biology and systems-level functional connectivity, a gap that matters clinically because the psychoplastogen literature and therapeutic-mechanism models point to durable circuit remodeling rather than acute receptor occupancy alone as the substrate of therapeutic benefit [1,3,4]. Future work should test concentration-response effects, longitudinal post-washout dynamics, and structural correlates of the network phenotype. Computational modeling and in vivo validation will be needed to determine whether this local network signature generalizes to intact circuits.

## Acknowledgements

We thank the Franck group at the University of Wisconsin-Madison for providing primary cortical neuron preparations.

## Code and data availability

The full analysis pipeline is available at https://github.com/hailab-uw/PsychoplastogensMEA. Raw TDT recording blocks are available from the corresponding author on reasonable request.

## Author contributions

Conceptualization: I.H., A.H., M.I.B.; Methodology: I.H., I.B., M.I.B., A.H.; Software: I.H., I.B., B.K.; Formal analysis: I.H.; Investigation: I.H., I.B., B.K., M.N.; Resources: I.B., A.H.; Data curation: I.H., I.B.; Writing - original draft: I.H.; Writing - review and editing: I.H., M.I.B., A.H., M.N., I.B., B.K.; Visualization: I.H., B.K.; Supervision: M.I.B., A.H.; Funding acquisition: A.H.

