## Supplementary material for "Acute DOI exposure drives cortical hyperexcitability and functional network remodeling": SI

### Contents

SM1 — MEA geometry, channel mapping, and spatial decay
SM2 — Connectivity estimation and STTC robustness
SM3 — Edge-density thresholding and density-sweep robustness
SM4 — Functional-network metrics and complete metric tables
SM5 — Hierarchical bootstrap procedure
SM6 — Supplementary visualization panels
SM7 — Supplementary visualization panels

### SM1 — MEA geometry, channel mapping, and spatial decay

#### Channel mapping and distance matrix

Recordings were performed on Multi Channel Systems 60MEA100/10iR chips with a standard 8 x 8 grid layout (four corner positions absent, yielding 60 MEA positions). On the iR model, one position (MCS label 15) is occupied by a large internal reference electrode, leaving 59 recording electrodes plus 1 reference. Inter-electrode pitch is 100 µm. The TDT amplifier headstage exposes 64 input channels; the 60 MEA positions map to PZ5 channels [1:15, 17:31, 33:47, 49:63], channel 15 carries the internal reference signal, and PZ5 channels 16, 32, 48, and 64 are grounded. Channel-to-grid mapping follows the MCS standard 60-electrode layout document, with the physical PZ5 channel index mapped to grid position (row, column) via the electrode-label look-up table.

The Cartesian coordinates of each electrode are derived from the grid position: x = column index, y = 9 - row index (so that row 1 plots at the top of the array). The pairwise Euclidean distance between electrodes i and j is:

$$d_{ij} = \sqrt{((x_i - x_j)^2 + (y_i - y_j)^2)}$$

where distances are expressed in electrode-pitch units (1 unit = 100 µm). The layout stores a 60 x 60 symmetric distance matrix over MEA positions, with analyses using the 59 x 59 recording-electrode subset after excluding the internal reference. Distances range from 1.0 (adjacent electrodes) to 9.90 (opposite corners of the active region). This distance matrix is used in the spatial-decay analysis (Figure 3f, Figure 5f), where edge weights are binned by inter-electrode distance in 1-pitch-unit intervals (0-1, 1-2, ..., 7-8) and summarized as median with interquartile range.

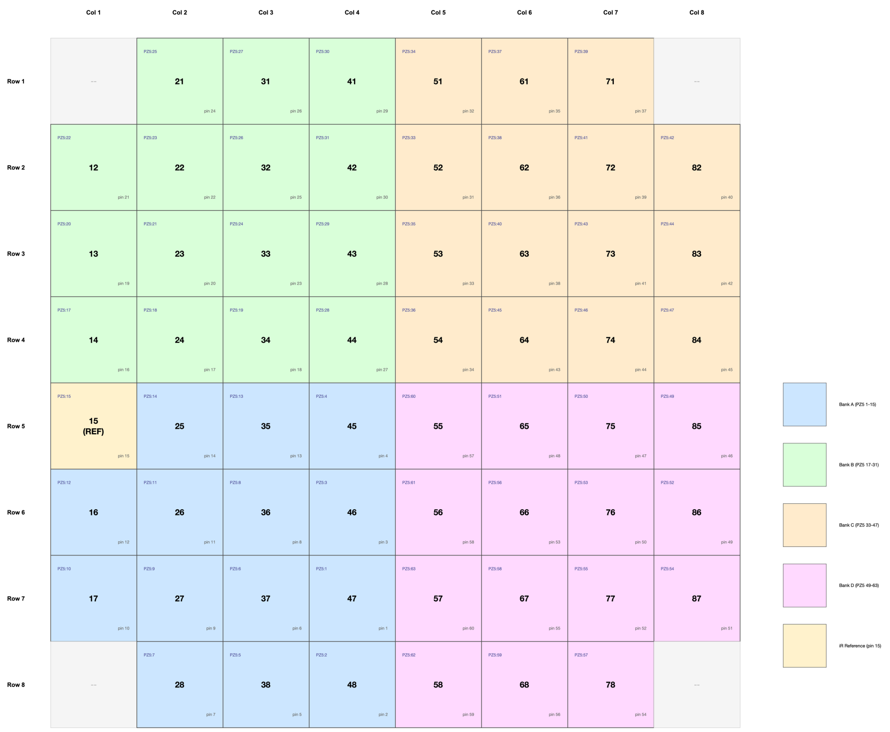


**Supplementary Figure S1.** *Complete channel mapping from TDT PZ5 digitizer indices to MCS electrode labels and 8 x 8 grid positions. Each cell shows the MCS electrode label (bold center), PZ5 channel index (top left), and MCS 1-dim pin number (bottom right). Cells are color-coded by PZ5 bank (A-D), and the internal reference electrode (MCS pin 15) is marked and excluded from the recording-electrode analysis set. This figure validates the channel-to-electrode correspondence used throughout the analysis pipeline.*

**Spatial-decay computation**

The spatial decay of functional connectivity (Figures 3f and 5f) was computed by binning all upper-triangular off-diagonal entries of the weighted (pre-binarization) adjacency matrix by inter-electrode Euclidean distance (above). Edge weights were pooled across all paired recordings within each arm, and the absolute value of the peak cross-correlation was used. For each distance bin (1-unit width), the median and interquartile range were computed across all pooled edges.

This visualization follows the link-length analysis of Downes et al. (2012) [1] (Figure 5), which demonstrated that functional connections in dissociated cortical MEA cultures exhibit a characteristic spatial decay, with nearby electrodes showing stronger correlations than distant pairs. Our spatial-decay panels show that DOI preserves this distance-dependence while redistributing coupling strength, and that the ketanserin arm shows a qualitatively similar pattern with reduced overall coupling magnitude.

To quantify whether treatment altered the shape (not just the magnitude) of the spatial decay, we computed a per-well ordinary least-squares regression slope of |edge weight| on inter-electrode distance for each recording. The paired Wilcoxon signed-rank test on per-well slopes showed no significant change for either the DOI arm ($p = 0.563$, $n = 6$) or the ketanserin arm (exact test, $p = 0.500$, $n = 3$), indicating that we did not detect a treatment-related change in distance-dependence under either pharmacological condition. These slope-test p-values are annotated on Figures 3f and 5f.

### SM2 — Connectivity estimation and STTC robustness

#### Cross-correlation connectivity estimation

Pairwise functional connectivity was estimated by binning each channel's spike train at 1 ms resolution, z-scoring the resulting count vector (subtracting the mean and dividing by the standard deviation), and computing the cross-correlation function over a +/-100 ms lag window using MATLAB's xcorr with 'unbiased' normalization. The peak value of the cross-correlation function within the lag window was taken as the connection weight, and the lag at which the peak occurred was retained as the peak-lag estimate. Self-connections (diagonal) were set to NaN.

The z-scoring step puts channels on a common scale before cross-correlation and helps make peak cross-correlation values comparable across channels with different firing rates. The 1 ms bin width was chosen to capture synaptic-timescale temporal relationships while maintaining computational tractability; the peak cross-correlation approach follows Garofalo et al. (2009) [2] and Poli, Pastore and Massobrio (2015) [3]. The +/-100 ms lag window captures both monosynaptic and polysynaptic delays in dissociated cortical cultures, where conduction velocities of 0.1-0.5 m/s across 100-800 μm inter-electrode distances yield expected delays of 0.2-8 ms.

#### STTC robustness analysis

The Pearson cross-correlation on z-scored binned spike counts is sensitive to absolute firing rate when rates differ between conditions (Cutts and Eglen (2014) [4], Table 3). To test whether the graph-metric effects reported in the main analysis persist under a rate-corrected connectivity estimator, we recomputed pairwise functional connectivity using the spike-time tiling coefficient (STTC) [4] with a synchrony half-window of +/-50 ms. STTC is bounded in [-1, 1] and measures the degree to which spikes from two trains co-occur within a temporal window, normalized by the fraction of recording time tiled by each train. It is designed to reduce firing-rate bias and is used here as the principal rate-control estimator.

From each STTC adjacency matrix, we derived the same six graph-theoretic metrics as in the main analysis, using the same 10 % edge-density thresholding. Supplementary Tables S5-S6 compare Pearson and STTC estimates. The goal is not to replace the main Pearson/10 % functional-network figures, but to bound the extent to which the Pearson graph metrics are rate-sensitive.

**Table S1.** *Pearson vs STTC comparison for the DOI arm (n = 6 wells).*

| Metric | Pearson Δ | STTC Δ | Same sign? |
|---|---|---|---|
| Edge density | 0.000 | 0.000 | Yes |
| Clustering coeff. | -0.015 | -0.011 | Yes |
| Char. path length | **-0.170** | **-0.100** | Yes |
| Small-worldness σ | +0.810 | -0.125 | No |
| Global efficiency | -0.020 | -0.019 | Yes |
| Modularity Q | **-0.051** | **-0.037** | Yes |

*Note. Median Δ is treatment minus baseline for each metric at 10 % edge density.*

The DOI path-length decrease persists under STTC but is weaker than under Pearson and directionally present in 4/6 wells rather than 5/6 wells. This supports path length as the most stable functional-network endpoint while bounding the strength of the Pearson-based claim. Modularity Q remains negative under both estimators. Small-worldness changes sign between estimators, reinforcing that functional-network family interpretation should emphasize the overall pattern and the more robust path-length axis rather than overinterpreting any single estimator-sensitive graph metric.

**Table S2.** *Pearson vs STTC comparison for the ketanserin arm (n = 3 wells).*

| Metric | Pearson Δ | STTC Δ | Same sign? |
|---|---|---|---|
| Edge density | 0.000 | 0.000 | Yes |
| Clustering coeff. | +0.062 | -0.047 | No |
| Char. path length | **-0.242** | **-0.468** | Yes |
| Small-worldness σ | +1.964 | +0.907 | Yes |
| Global efficiency | -0.037 | -0.085 | Yes |
| Modularity Q | **+0.040** | **-0.069** | No |

*Note. Same format as Table S1. This arm is reported descriptively.*

Path-length shortening is present under both estimators in the ketanserin arm, but because n = 3 this result is descriptive. The modularity sign reversal (Pearson: +0.040 vs STTC: -0.069) indicates that the small Pearson-based modularity signal in the ketanserin arm is estimator-sensitive and should not be interpreted as confirmatory.

### SM3 — Edge-density thresholding and density-sweep robustness

#### Edge-density thresholding and binarization

Adjacency matrices were binarized at a target edge density of 10 % using proportional thresholding as described by Rubinov and Sporns (2010) [5]. For each adjacency matrix, the threshold was set from the upper-triangular off-diagonal entries so that the strongest 10 % of possible edges survived thresholding. This data-driven approach avoids the need for an absolute threshold, which would be sensitive to recording duration and bin width.

The 10 % density was chosen as the primary analysis value because (a) it lies within the 5-20 % range recommended by Rubinov and Sporns (2010) [5] for sparse biological networks, (b) it yields connected graphs for all recordings in both arms, and (c) the Downes et al. (2012) [1] developmental MEA study used a comparable density range. Robustness to this choice was tested across a density sweep of 5, 10, 15, and 20 % edge density (density-sweep analysis below).

#### Density-sweep robustness analysis

To test the sensitivity of graph-theoretic results to the binarization threshold, network metrics were recomputed at target edge densities of 5, 10, 15, and 20 %, consistent with the sparse-network range recommended by Rubinov and Sporns (2010) [5]. Supplementary Tables S3 and S4 report median baseline-to-treatment changes for characteristic path length, small-worldness σ, and modularity Q across the density sweep. These analyses are complementary to the primary 10 % density functional-network endpoints shown in the main text.

**Table S3.** *Density sweep for the DOI arm (n = 6 wells).*

| Target density | Median Δ path length | Median Δ small-worldness σ | Median Δ modularity Q |
|---|---|---|---|
| 5 % | +0.060 | -1.356 | -0.038 |
| 10 % (primary) | **-0.170** | **+0.810** | **-0.051** |
| 15 % | -0.182 | +0.602 | -0.044 |
| 20 % | -0.002 | +0.295 | -0.039 |

*Note. Median Δ is reported for characteristic path length, small-worldness σ, and modularity Q at each target edge density.*

At the primary 10 % density, the DOI arm shows the same pattern as the main analysis: shorter path length, higher small-worldness σ, and lower modularity Q. Across the 5-20 % sweep, modularity Q trends negative throughout, whereas path length and small-worldness are density-sensitive at the sparsest 5 % threshold and should be interpreted around the prespecified 10 % density rather than as threshold-invariant standalone claims.

**Table S4.** *Density sweep for the ketanserin arm (n = 3 wells).*

| Target density | Median Δ path length | Median Δ small-worldness σ | Median Δ modularity Q |
|---|---|---|---|
| 5 % | +0.023 | -1.893 | +0.048 |
| 10 % (primary) | **-0.242** | **+1.964** | **+0.040** |
| 15 % | +0.220 | +0.420 | +0.074 |
| 20 % | +0.066 | +0.509 | +0.071 |

*Note. Same format as Table S3. This arm is reported descriptively.*

The ketanserin arm is threshold-sensitive and underpowered for confirmatory graph-metric inference. At the primary 10 % density, path length decreases and small-worldness σ increases, but the signs across the density sweep are not stable. Modularity Q is near zero or positive under Pearson across the tested densities, reinforcing the decision not to treat ketanserin modularity as confirmatory.

### SM4 — Functional-network metrics and complete metric tables

#### Complementary metric definitions

The main text defines the prespecified functional-network endpoints. The complementary metrics reported in Supplementary Tables S5-S6 are defined here so that the complete connectivity output is interpretable without repeating main-text equations.

**Edge density.** The fraction of possible edges present in the binarized graph. In our design, density is fixed at 10 % by construction:

$$\text{density} = \frac{2E}{\text{N}(\text{N} - 1)}$$

where *E* is the number of edges and *N* is the number of nodes.

**Clustering coefficient.** The mean local clustering coefficient across all nodes with degree >= 2. For each node *i*, the local clustering coefficient is the fraction of triangles formed by its neighbors relative to the maximum possible:

$$C_i = \frac{2t_i}{k_i(k_i - 1)}$$

where $t_i$ is the number of triangles through node i and $k_i$ is its degree [6].

**Global efficiency.** The average inverse shortest-path distance. Unlike characteristic path length, global efficiency handles disconnected components naturally because $1/\infty = 0$ [7]:

$$E_{glob} = \frac{1}{\text{N}(\text{N} - 1)} \Sigma \frac{1}{d_{ij}}$$

#### Complete connectivity metrics

**Table S5.** *Complete network-metric summary for the DOI arm (n = 6 wells).*

| Metric | Median baseline | Median treatment | Δ | 95% CI | Bootstrap BH p | Wilcoxon BH p | Hedges g (average-SD) |
|---|---|---|---|---|---|---|---|
| Edge density | 0.100 | 0.100 | 0.000 | [0.000, 0.000] | NA | NA | 0.00 |
| Clustering coeff. | 0.387 | 0.356 | -0.015 | [-0.073, +0.093] | NA | NA | +0.02 |
| Char. path length | 1.708 | 1.391 | -0.170 | [-0.676, -0.069] | <0.001 | 0.031 | -0.93 |
| Small-worldness σ | 5.684 | 6.458 | +0.810 | [+0.090, +2.270] | 0.022 | 0.031 | +1.18 |
| Global efficiency | 0.165 | 0.132 | -0.020 | [-0.163, +0.048] | NA | NA | -0.51 |
| Modularity Q | 0.094 | 0.054 | -0.051 | [-0.179, -0.004] | 0.022 | 0.031 | -0.40 |

*Note. For each graph-theoretic metric: median baseline, median treatment, median Δ, 95 % bootstrap CI of Δ, bootstrap BH-adjusted p where applicable, Wilcoxon BH-adjusted p where applicable, and Hedges' g (average-SD). The primary functional-network family is characteristic path length, modularity Q, and small-worldness σ; density, clustering coefficient, and global efficiency are complementary metrics. Endpoint interpretation follows the Wilcoxon BH-adjusted p column for the prespecified topology family.*

**Table S6.** *Complete network-metric summary for the ketanserin arm (n = 3 wells).*

| Metric | Median baseline | Median treatment | Δ | 95% CI | Bootstrap BH p | Wilcoxon BH p | Hedges g (average-SD) |
|---|---|---|---|---|---|---|---|
| Edge density | 0.100 | 0.100 | 0.000 | [0.000, 0.000] | NA | NA | 0.00 |
| Clustering coeff. | 0.420 | 0.476 | +0.062 | [-0.018, +0.083] | NA | NA | +0.61 |
| Char. path length | 1.714 | 1.471 | -0.242 | [-0.431, -0.181] | <0.001 | 0.375 | -0.53 |
| Small-worldness σ | 5.335 | 7.606 | +1.964 | [+0.772, +2.271] | <0.001 | 0.375 | +0.65 |
| Global efficiency | 0.187 | 0.150 | -0.037 | [-0.123, +0.009] | NA | NA | -0.54 |
| Modularity Q | 0.204 | 0.331 | +0.040 | [-0.167, +0.277] | 0.527 | 0.750 | +0.20 |

*Note. Same format as Table S5. Because n = 3, these graph metrics are reported descriptively, and inferential claims are not made from ketanserin p-values.*

**SM5 — Hierarchical bootstrap procedure**

The two-level hierarchical bootstrap (Saravanan et al. (2020) [8]) is used as a supplementary sensitivity analysis for the nested structure of MEA data (electrodes within wells). Exact Wilcoxon signed-rank tests on per-well summaries remain the primary inference because wells, not electrodes, are the independent experimental units. At each of B = 10,000 iterations:

1. Resample n recordings (wells) with replacement from the n available.

2. Within each resampled recording, resample electrodes with replacement.

3. Pool resampled electrodes and compute the median percent change.

The 95 % confidence interval is the [2.5, 97.5] percentile interval of the B bootstrap replicates. The two-sided bootstrap sensitivity p-value is 2 x min(fraction of replicates <= 0, fraction of replicates >= 0), capped at 1. The random seed is fixed at 1 for reproducibility. These bootstrap p-values are not used as the primary significance criterion.

### SM6 — Supplementary visualization panels

#### Per-recording exemplar panels

To enable independent visual inspection of every recording pair, exemplar panels (spike rasters and cross-correlation connectivity maps) were generated for all 6 DOI pairs and all 3 ketanserin pairs. In the main text, Figures 2 and 4 show panels from a single representative pair; the supplementary figures below present the equivalent panels for every recording pair in each arm. All panels were generated with the same pipeline parameters as the main-text figures (bandpass 300-2500 Hz, spike threshold 5.0 SD, cross-correlation bin 1 ms, +/-100 ms lag window, 10 % edge density).

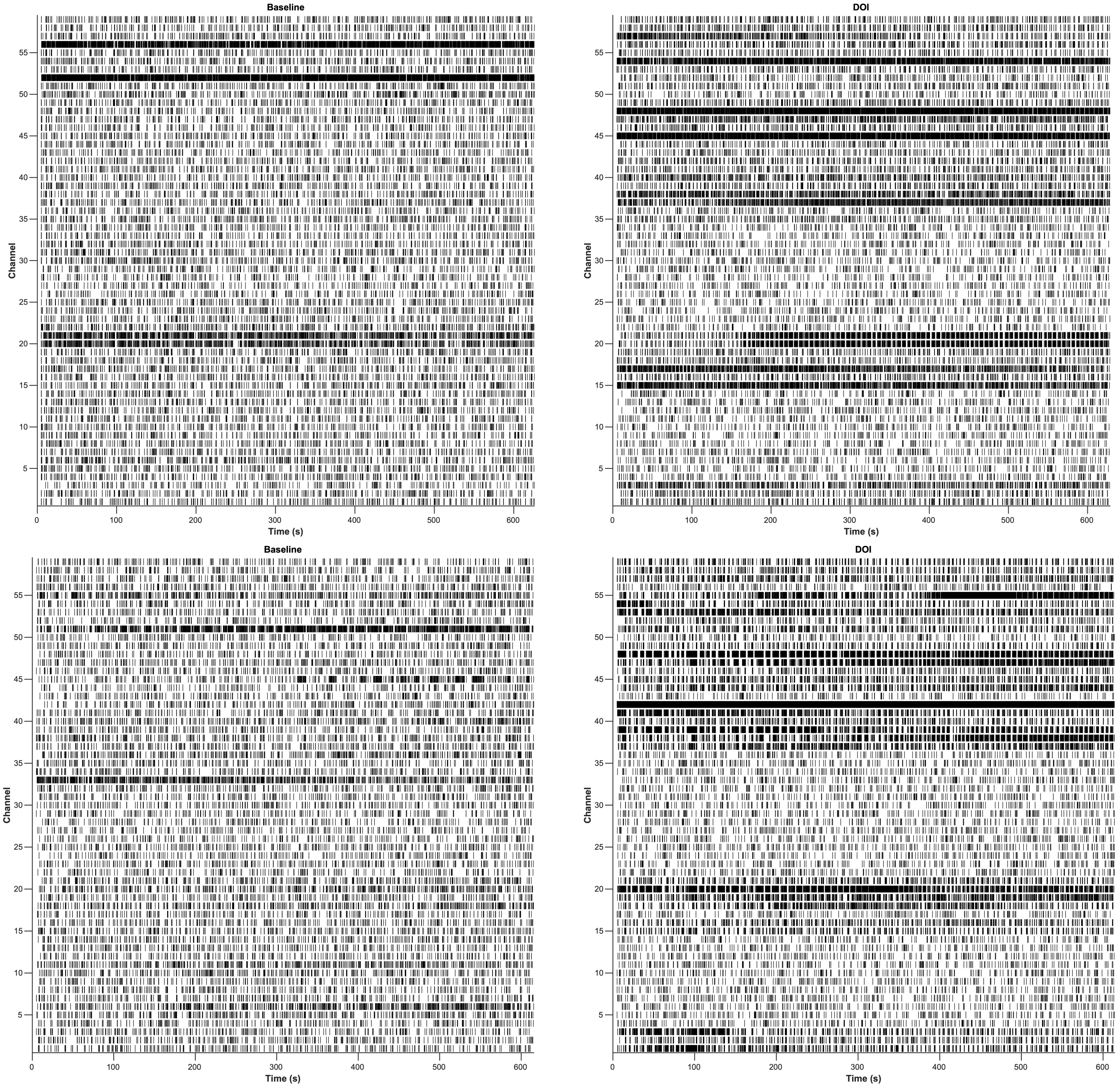

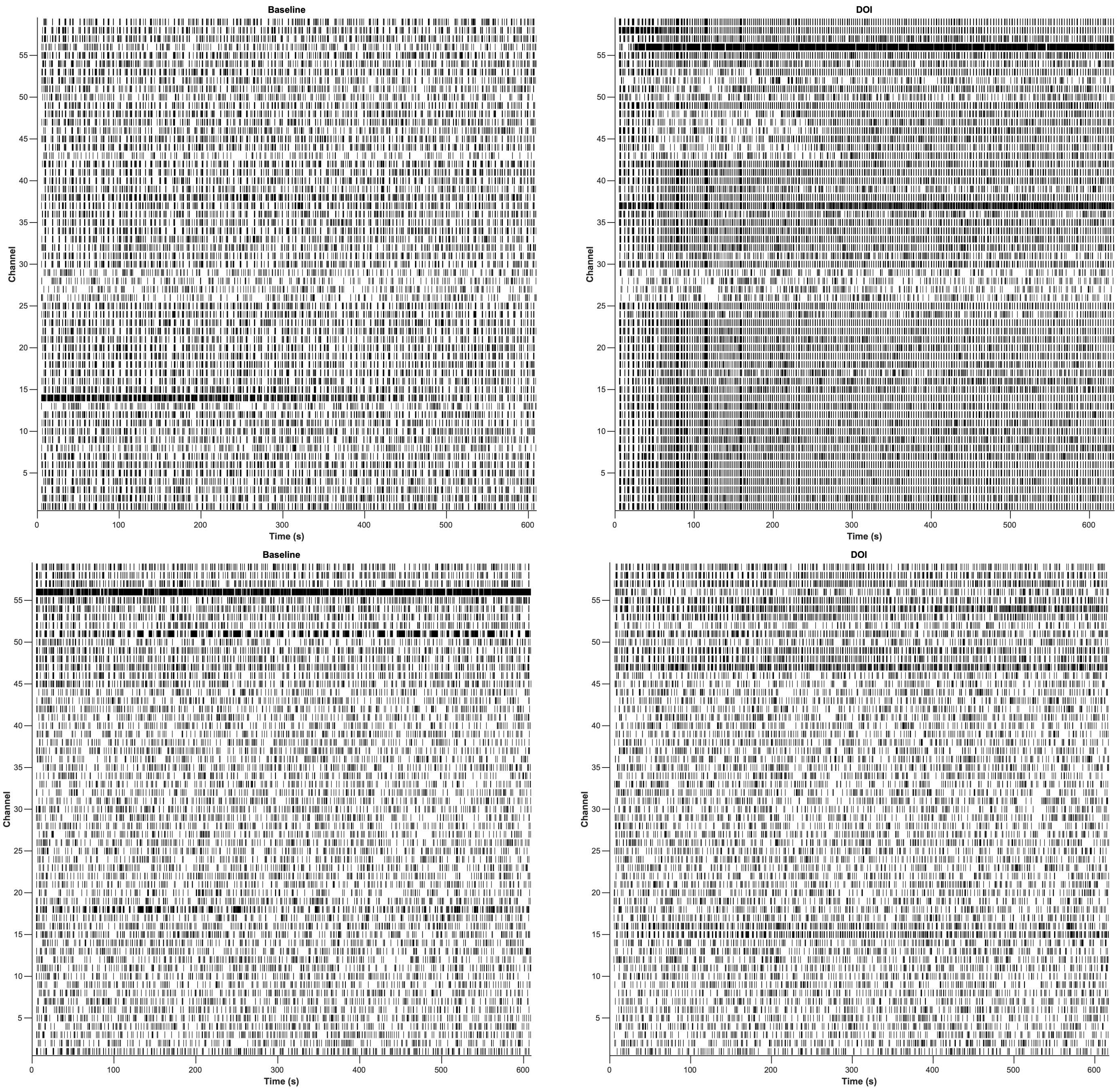

Baseline
DOI
Baseline
DOI
Channel
Time (s)

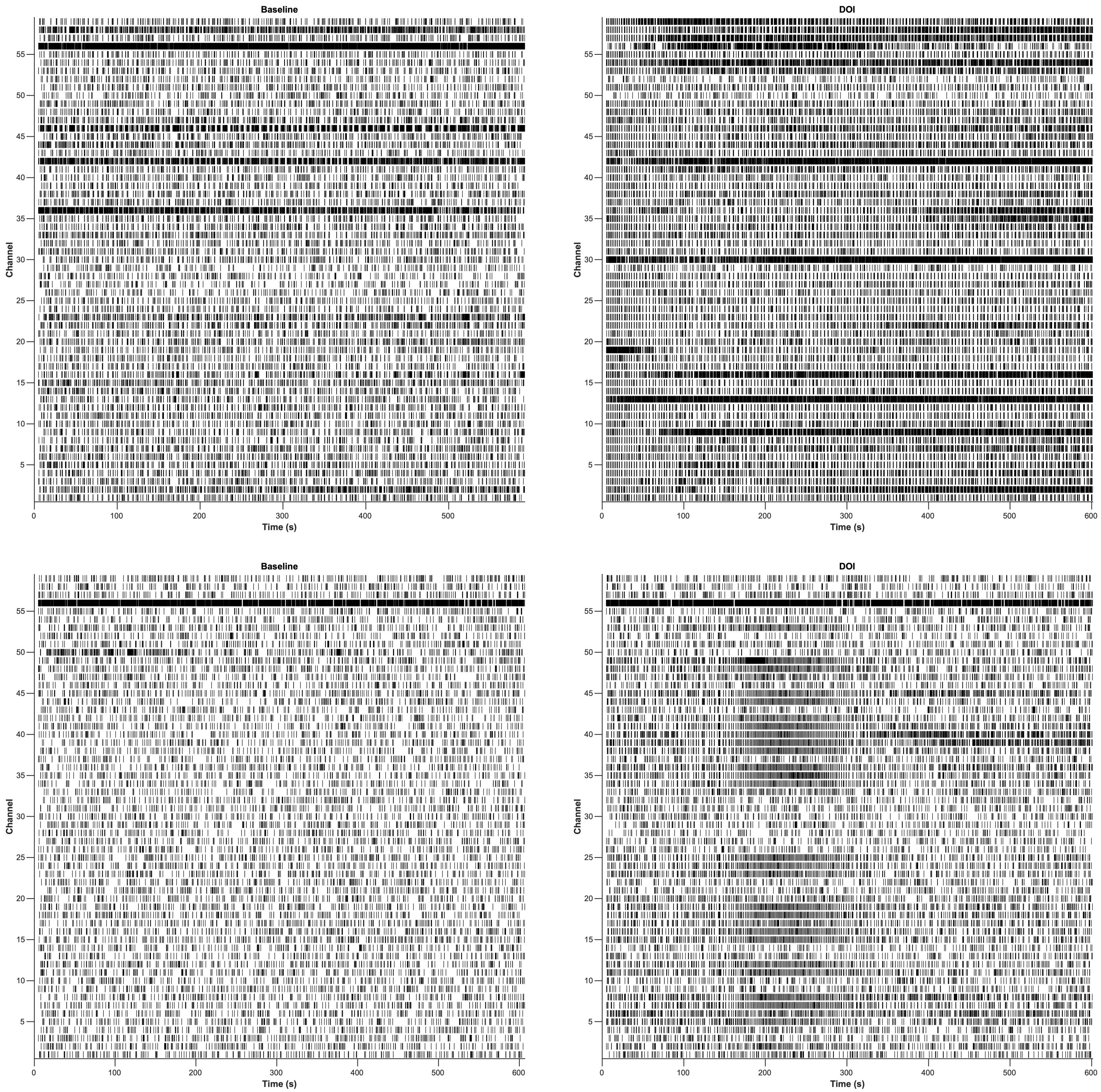


**Supplementary Figure S2.** *Per-recording spike rasters for all DOI pairs (pair 1-6). Each panel shows the full-recording spike-time raster for all 60 MEA channels. Left column: baseline; right column: post-acute DOI. Spikes are drawn as vertical ticks at their detection time.*

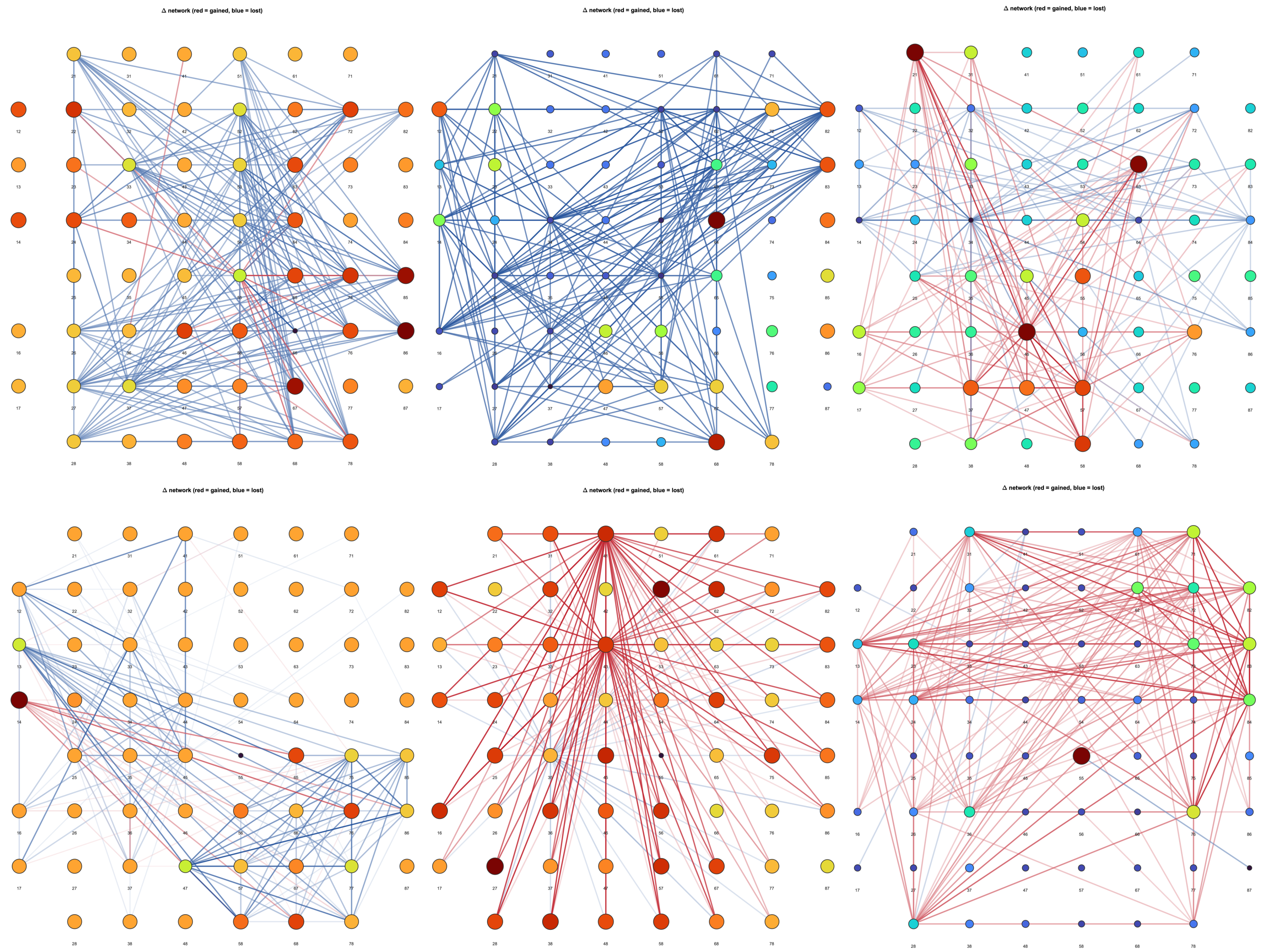


**Supplementary Figure S3.** *Per-recording functional connectivity difference networks for all DOI pairs (pair 1-6). Each panel shows post-exposure-minus-baseline network edges on the recording-electrode MEA geometry; edge color and width encode the signed change in connection weight, and labels identify the recording pair.*

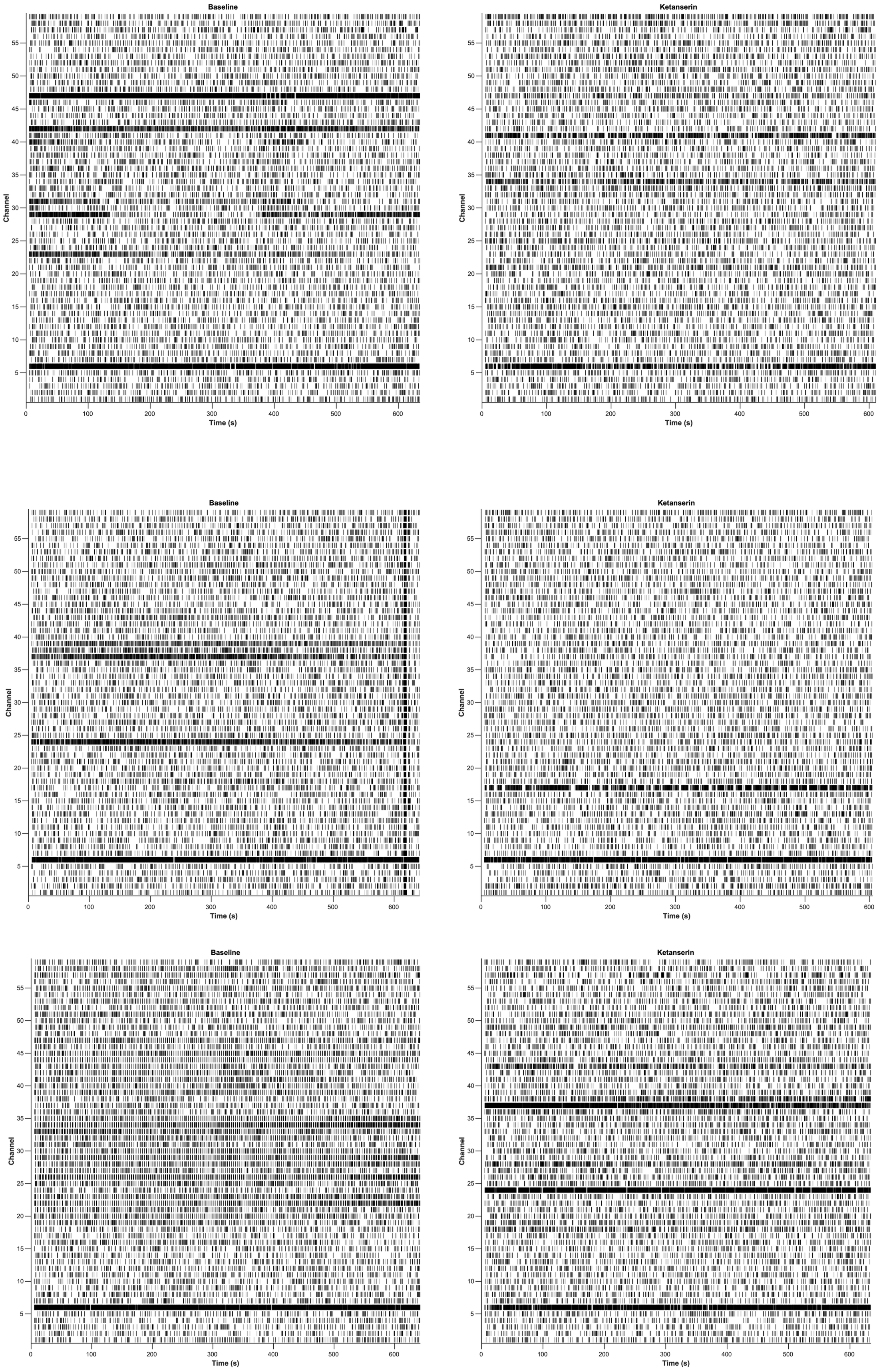


**Supplementary Figure S4.** *Per-recording spike rasters for all ketanserin pairs (pair 1-3). Same format as Supplementary Figure S2.*

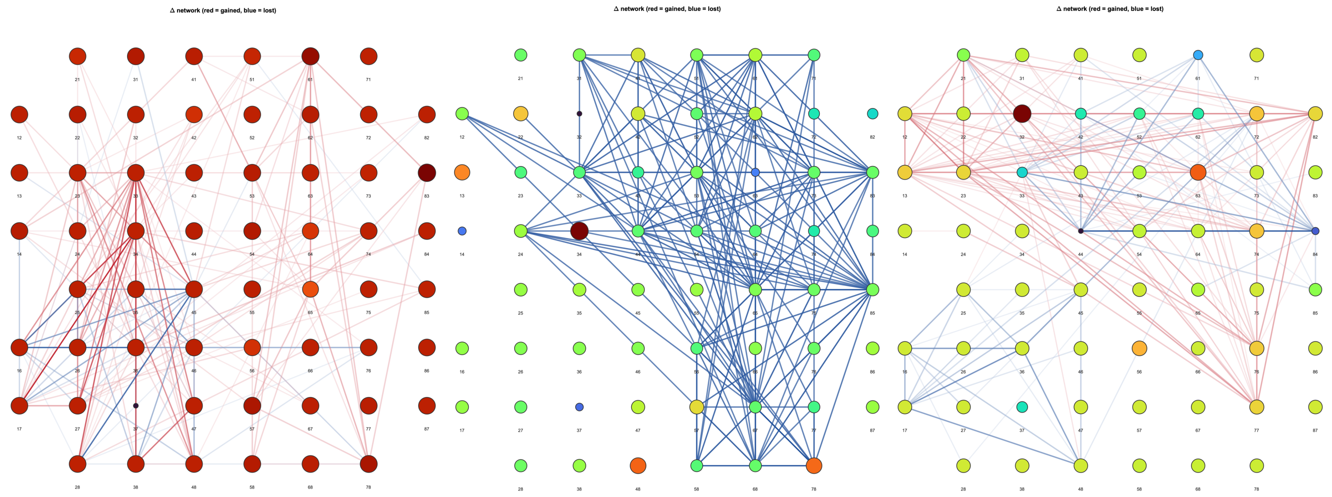


**Supplementary Figure S5.** *Per-recording functional connectivity difference networks for all ketanserin pairs (pair 1-3). Same format as Supplementary Figure S3.*

**Burst dynamics supplementary panels**

To enable detailed assessment of burst-level activity at finer temporal resolution than the full-recording rasters shown in the main text, two additional burst-focused visualization types were generated for both experimental arms.

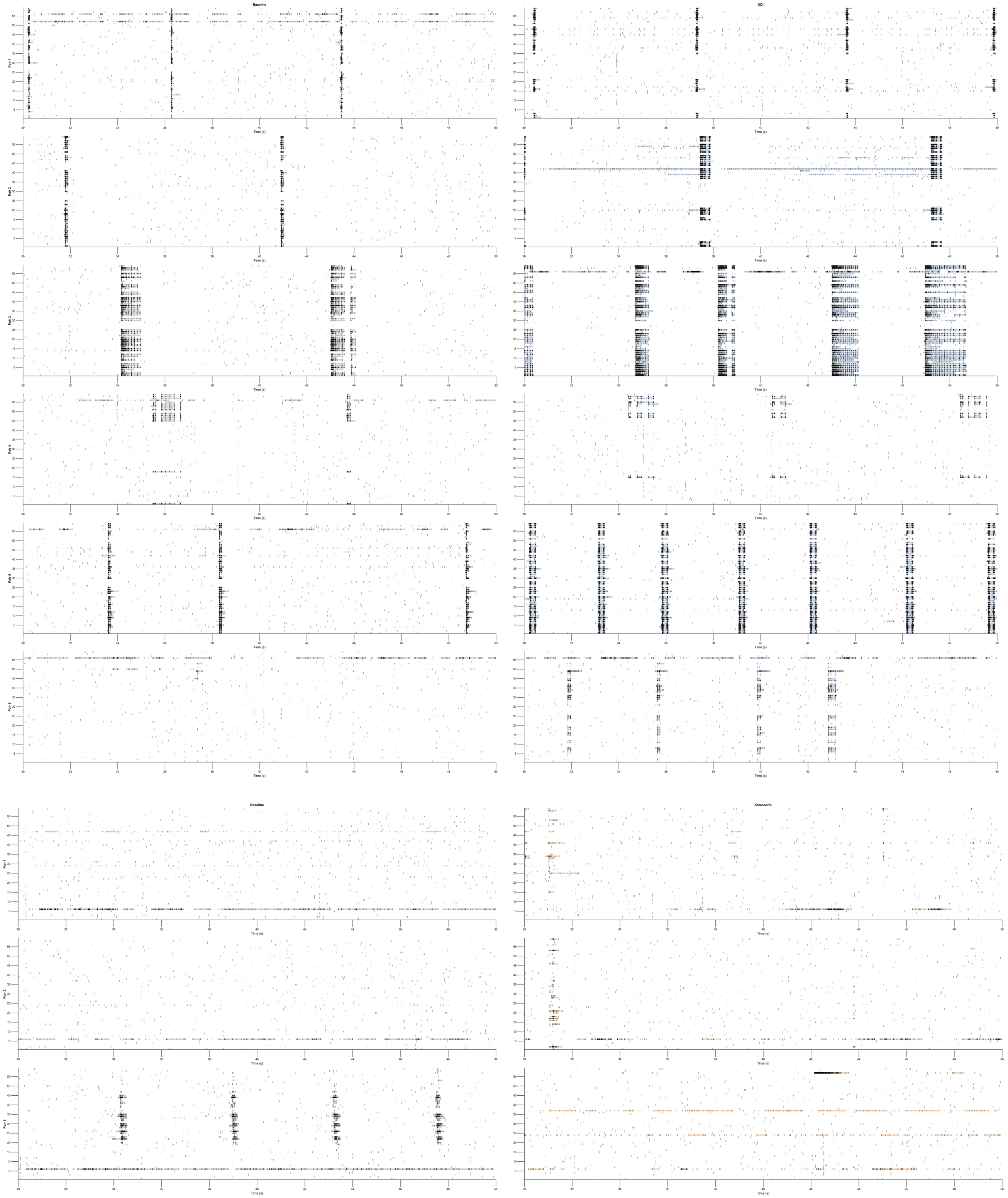


**Supplementary Figure S6.** *Zoomed 20-second spike rasters with burst-interval shading for all pairs. Each subplot shows a 20-second window (t = 30-50 s) of the spike raster for all 59 recording electrodes, with detected burst intervals shaded in translucent color behind the spike ticks. Left column: baseline; right column: treatment. This temporal resolution allows individual burst events (lasting 50-200 ms) to be visually resolved, whereas full-recording rasters compress bursts into single vertical lines.*

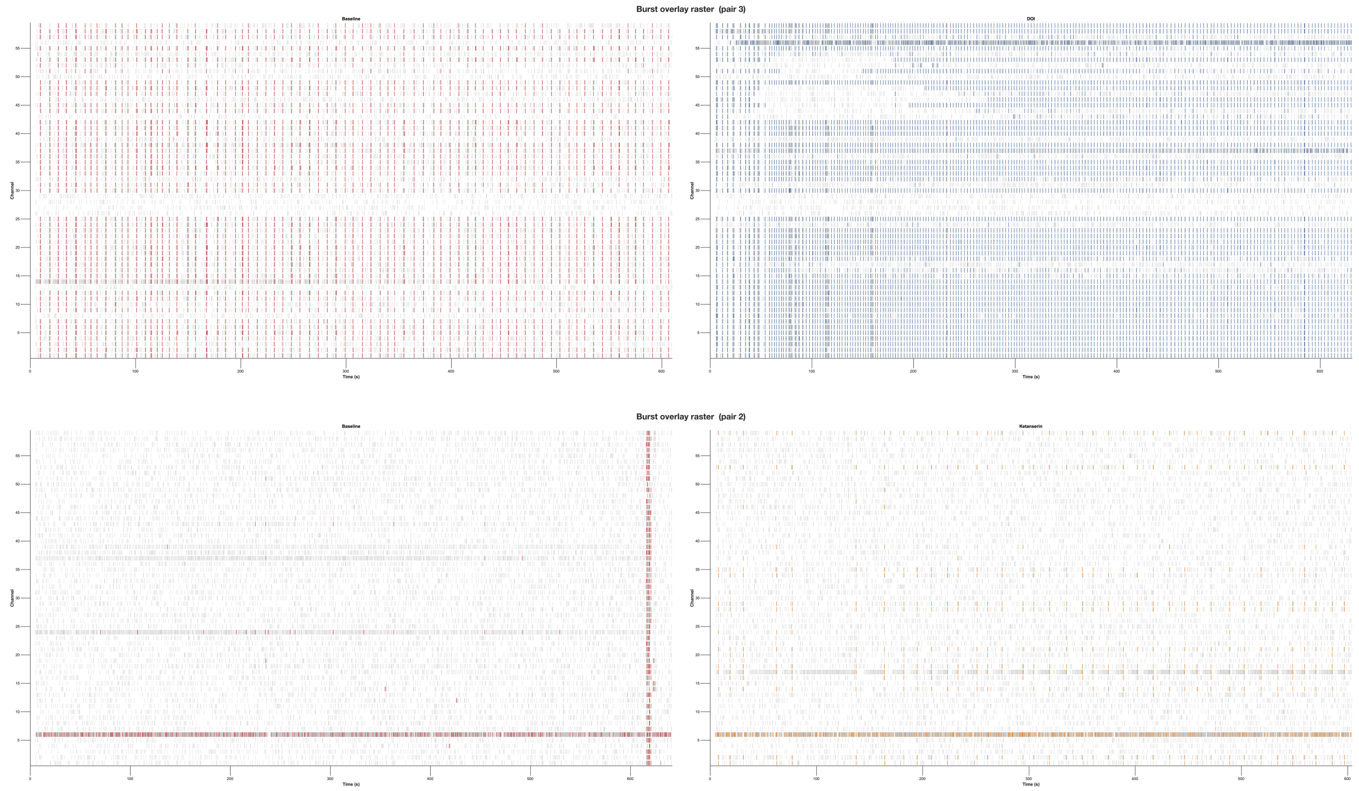


**Supplementary Figure S7.** *Full-recording rasters with burst-onset markers for one representative pair per arm (DOI pair 3, ketanserin pair 2). Tonic spikes are drawn in light gray; burst onset times are marked as colored vertical ticks, distinguishing burst-associated activity from tonic firing. This visualization separates the two activity modes that contribute differently to functional connectivity.*

**SM7 — Population firing-rate temporal stability**

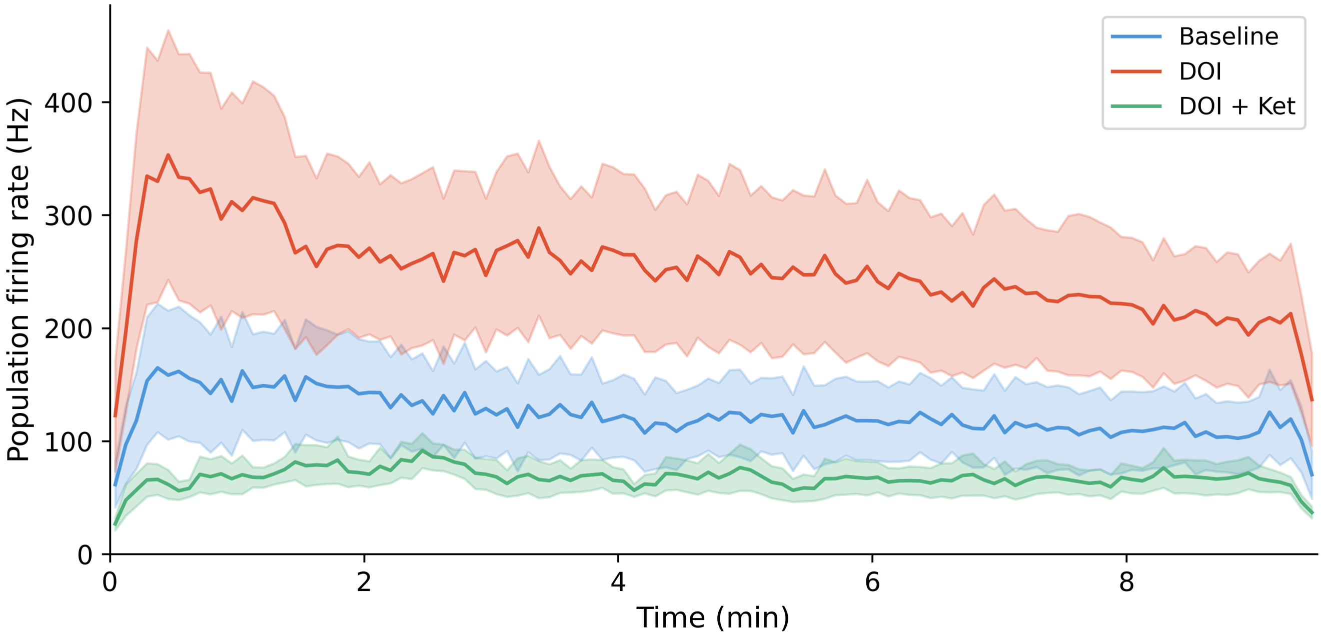


**Supplementary Figure S8.** *Population firing rate over the full shared recording duration (~9.5 min) for all three conditions: baseline (blue), post-acute DOI (red), and post-acute DOI + ketanserin (green). Each well's spike times were pooled across all channels into a single population spike train, binned at 5 s resolution, and smoothed with a 5-bin moving average. Lines show the mean across wells; shaded ribbons show +/- 1 standard deviation. All three conditions exhibit a brief transient increase in the first ~30 s, after which firing rate stabilizes. The relative separation between conditions (DOI > baseline > ketanserin) is maintained throughout, indicating that the rate effects reported in the main text are not driven by initial perturbation artifacts.*
